# Best practices to cluster large molecular libraries

**DOI:** 10.1101/2025.11.28.691214

**Authors:** Kenneth Lopez-Perez, Ramon Alain Miranda-Quintana

## Abstract

BitBIRCH is a novel clustering algorithm that enables the analysis of extremely large molecular libraries; however, its performance can be hindered by an excessive number of singletons or the formation of disproportionately large clusters. Here, we present a data-driven strategy to identify optimal BitBIRCH parameters that mitigate these limitations. Using the ChEMBL34 library as a case study (with additional datasets reported in the Supporting Information), we show that similarity thresholds between three and four standard deviations above the global mean provide a balanced trade-off between cluster count and medoid similarity. These values are efficiently approximate with the iSIM and iSIM-sigma frameworks. For the branching factor, values as high as computationally feasible are recommended, as increasing it to 1024 substantially reduced the number of singletons. We further introduce an iterative re-clustering procedure wherein the similarity threshold can be adjusted to merge related subclusters and singletons from the initial clustering, providing user-defined control over the extent of cluster fusion. This work provides practical guidelines to enhance the robustness and usability of BitBIRCH for large-scale molecular clustering.

## Communication

The recently introduced BitBIRCH^1^ algorithm leverages a hierarchical tree structure and the Instant Similarity (iSIM)^2^ formalism to efficiently cluster molecules represented by binary fingerprints. BitBIRCH enables the clustering of ultra-large molecular libraries with unprecedented scalability based on Tanimoto similarity^3,4^. With recent advances in memory management and computational efficiency, it can now cluster one billion molecules in approximately two hours.^5^ In the original publication^1^, the quality of the clusters produced by BitBIRCH was evaluated on smaller datasets and found to be comparable to those obtained with classical similarity-based algorithms such as Taylor–Butina^6,7^. However, when applied to larger datasets, challenges in cluster quality have been observed. Most notably, the emergence of a loosely defined, highly populated cluster and an excessive number of singletons^8^, issues also encountered with the Taylor–Butina algorithm^9^ (or any other sphere-based method). The former issue has been previously addressed through refinement strategies designed to subdivide overly large clusters.^8^ In this communication, we present a set of practical recommendations and straightforward strategies to improve BitBIRCH clustering outcomes. Specifically, we provide guidance for selecting appropriate similarity thresholds and incorporating singletons into clusters. These practices are particularly relevant for large-scale datasets, where direct assessment of clustering quality is more challenging, thus offering new BitBIRCH users a clearer understanding of how to optimize the algorithm for their specific applications.

Some studies have attempted to define what values can define a molecule as “similar” in the eyes of the similarity principle^10^, our analyses indicate that such a value is inherently dataset- and fingerprint-dependent within the context of a set. The overall sparsity of a fingerprint strongly influences the distribution of Tanimoto similarity values: sparse fingerprints typically yield lower similarity distributions compared to more densely populated ones. In the past this has been pointed out when comparing distributions between MACCS^11^ and ECFP4^12,13^ fingerprints.^10^ **Figure 1a)** compares the pairwise Tanimoto similarity distributions for a carotenoid dataset (*n* = 1038) from COCONUT^14^ represented with ECFP4 and RDKit^15^ fingerprints. The average similarity differs by approximately 0.1 between the two representations, with ECFP4 exhibiting greater positive kurtosis than RDKit. Clearly, using a fixed threshold in an algorithm that relies on average cluster would lead to variant results across different representations. **Figure 1b)** shows the similarity distributions for the same carotenoid set and for a random sample of the same size from ChEMBL34^16^, both represented with ECFP4 fingerprints.

**Figure 1.**
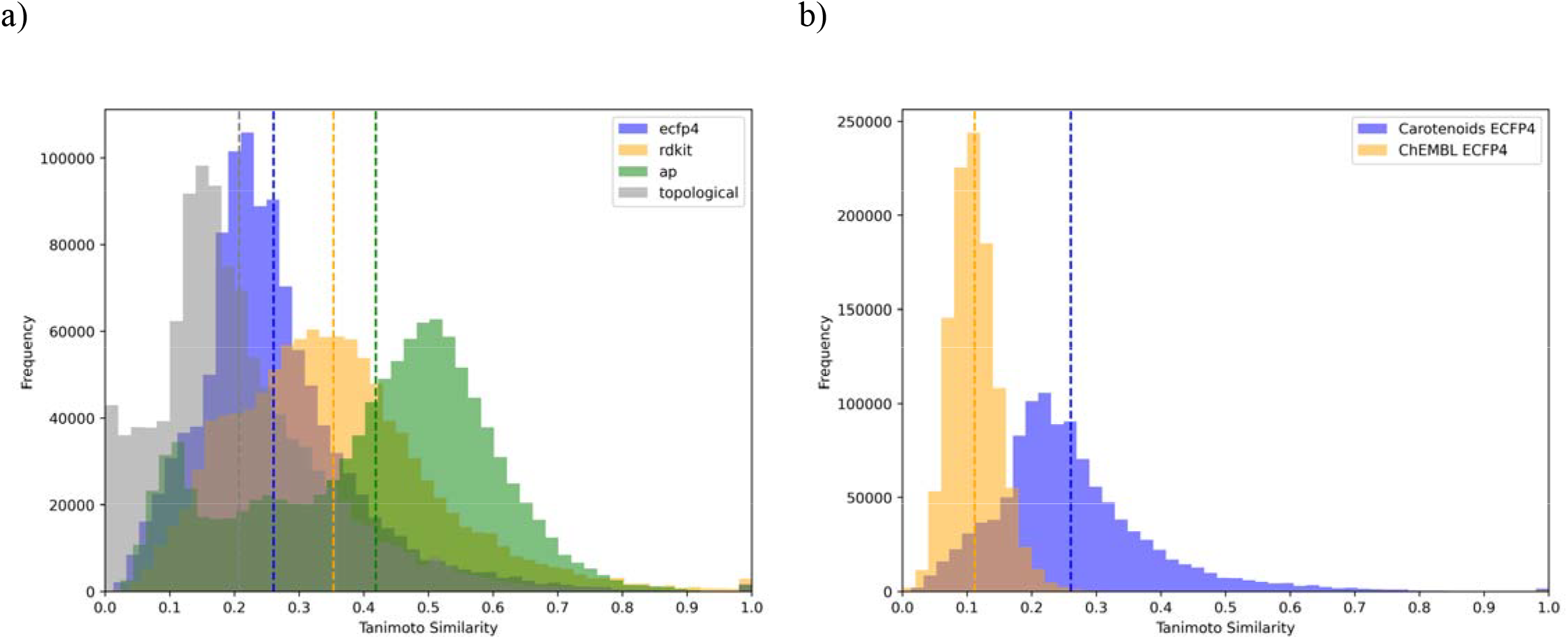
Pairwise Tanimoto similarity distributions for a) carotenoids from the COCONUT library represented with ECFP4 and RDKIT binary fingerprints, b) the carotenoid COCONUT set and a random sample of the same size from ChEMBL34.

As expected, the random sample exhibits a lower average similarity than the carotenoid-specific dataset, along with a distinct difference in kurtosis. These results demonstrate that applying a single similarity threshold across databases can lead to poor clustering performance: low thresholds produce a few heterogeneous clusters, while high thresholds generate excessive singletons and few meaningful clusters. Based on these observations, we propose an informed threshold selection strategy that leverages the average similarity and standard deviation of pairwise similarities to adapt the threshold to each dataset.

However, because the primary goal of BitBIRCH is to avoid explicit pairwise similarity calculations, it would be counterproductive to compute the full pairwise matrix merely to define a threshold. Instead, we can estimate them efficiently and with linear complexity using the iSIM^2^ and iSIM-sigma^17^ formalisms, which approximate the average similarity and the standard deviation with high accuracy, with minimal computational overhead.

**Figure 2a)** shows the number of clusters and singletons obtained when clustering the complete ChEMBL34 library (excluding compounds flagged as inorganic, *n* = 2,764,449) using different similarity thresholds. Each step in the plot corresponds to an increase of one standard deviation above the average similarity. As expected, the number of singletons converges with the total number of clusters as the threshold approaches 1. This plot provides a practical reference for anticipating the number of clusters generated at different thresholds. For a dataset of this size, thresholds corresponding to three to four standard deviations away from the mean similarity have a reasonable level of data reduction suitable for most applications. Thresholds below this range (e.g., 0-20 standard deviations) tend to produce overly large and diffuse clusters, as illustrated by the emergence of clusters with over dominant populations for most fingerprint types (see Figure S2). Thresholds corresponding to more than 5 standard deviations have over 1 million clusters, and a singleton/cluster ratio above 0.50; these numbers suggest that beyond 5 standard deviations there is an over-partition that is not practical for most applications.

**Figure 2.**
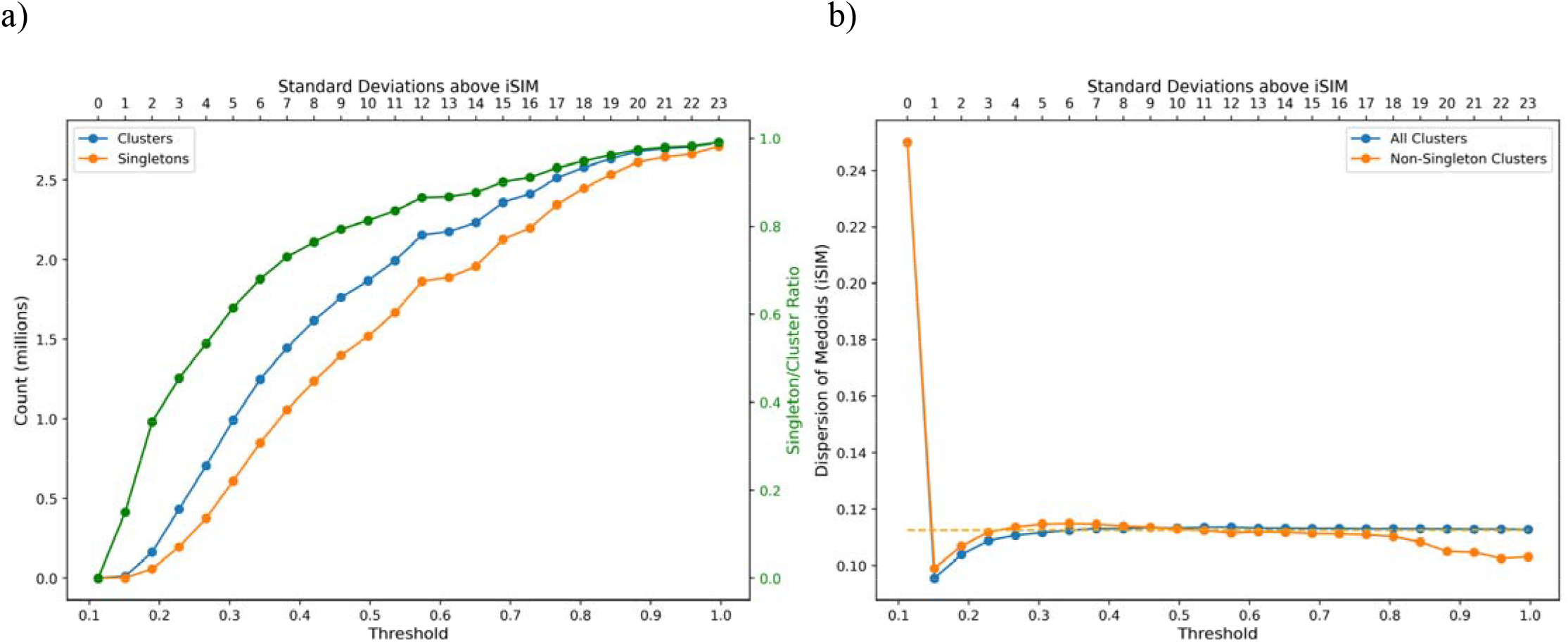
Clustering results for ChEMBL34 represented with 2048-bit ECFP4 binary fingerprints using BitBIRCH with diameter merging and a branching factor of 50. a) Variation of the number of clusters, singletons, and singleton/cluster ratio with respect to the threshold. b) iSIM values of the cluster medoids. The horizontal dashed line indicates the iSIM of the complete dataset. The top axis represents the iSIM-sigmas added to the iSIM of the whole set corresponding to each threshold.

A similar analysis performed with other fingerprints (see Figure S1) produced consistent clustering behavior within this range, despite their different absolute threshold values. Our observations are consistent with previously reported threshold choices for Taylor–Butina clustering. A prior study proposed 0.30 as a general similarity threshold for ECFP-based fingerprints^18^; that work evaluated only increments of 0.1 on a random subset of ZINC. Our approach suggests thresholds of approximately 0.27 (four standard deviations) for a random sample from ChEMBL, value that closely aligns with the earlier recommendation. Similarly, a separate clustering study on a cancer-focused library using ECFP4 fingerprints identified 0.90 as the most suitable threshold.^19^ The average library similarity of this particular library, after outlier removal, was greater than 0.70. This is consistent with our findings, probably after adding 3 or 4 standard deviations the resulting value would be close to 0.90. Taken together, our results and their agreement with previous reports underscore that no universal similarity cutoff exists across fingerprint types or databases. Instead, a data-driven strategy grounded in the distribution of similarities provides a more robust and generalizable basis for selecting clustering thresholds.

Beyond the number of clusters and singletons, we propose the usage of the iSIM of the medoids as a metric to determine if the clusters have a partition that resembles the chemical space of the whole set. In **Figure 2b)** we show the variation of the iSIM of the medoids at different thresholds in increasing steps in standard deviation. As expected at larger thresholds the iSIM of the medoids will tend to the iSIM of the whole set, this is because at those large thresholds most of the clusters are singletons hence it would be close to calculating the iSIM of the individual fingerprints. To make sure that the partition is good, we include the iSIM of the medoids excluding singleton clusters. For most applications what we want is the least number of clusters that are representative of the whole set, we can see that in **Figure 2b)** we get close to the iSIM of the whole set with 3 to 4 standard deviations above the iSIM of the set. These observations remain for other types of fingerprints as shown on Figure S1.

Up to this point, we have examined the influence of the similarity threshold alone; however, BitBIRCH includes a second key parameter, the branching factor, which has thus far been kept at its default value of 50. Although selecting the threshold based on iSIM statistics alleviates the excessive formation of singletons, their proportion remains above 0.4 even when using proposed optimal threshold range iSIM. Due to the tree structure of BitBIRCH, for large dataset molecules or subclusters may traverse branches without meeting similar counterparts, resulting in residual singletons or small clusters. Adjusting the branching factor directly controls the maximum number of subclusters or molecules to which an incoming molecule is compared at each tree level, and therefore has a direct impact on clustering compactness and singleton prevalence. In Figure 3, we investigate the effect of varying the branching factor on both the total number of clusters and the iSIM of the resulting medoids, while maintaining the threshold determined in the previous section (three to four standard deviations above the global iSIM of the dataset).

**Figure 3.**
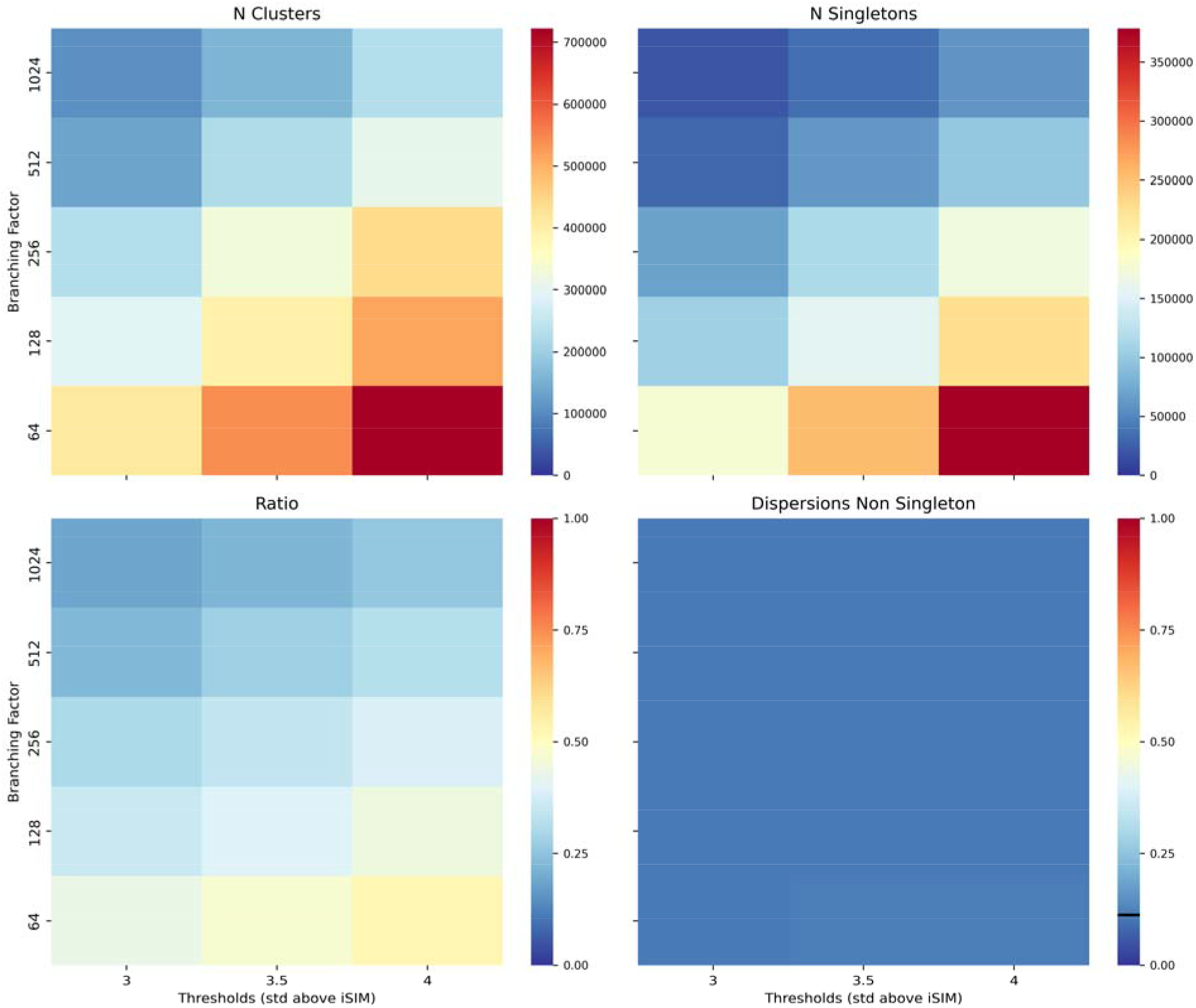
Variation of the number of clusters, number of singletons, ratio singleton/clusters and the dispersion (iSIM of medoids) of non-singleton clusters with respect to the branching factor and the thresholds, for the ChEMBL34 library represented with 2048-bit binary ECFP4 fingerprints. iSIM of the whole library is marked with a black line in the dispersions panel.

A first notable observation is that the iSIM of the medoids remains close to the iSIM of the entire dataset across all tested combinations of thresholds and branching factors, indicating that the selected threshold range (three to four standard deviations above the global iSIM) is indeed appropriate independently of the branching factor. As expected, increasing the branching factor leads to a reduction in the number of singletons, confirming that higher branching values help mitigate excessive singleton formation. These trends are consistent across the other fingerprint types evaluated (see Figures S3-S6). Nevertheless, for Atom Pair and RDKit fingerprints, although a substantial reduction in singletons is observed, the singleton-to-cluster ratio remains comparatively high, suggesting that these representations are inherently more prone to fragmenting into smaller clusters under these conditions.

An analysis of the computational performance associated with different thresholds and branching factors is presented in Figure 4. The results show that clustering time increases with the branching factor; however, this trend exhibits a semi-plateau behavior, and the largest observed difference across the tested values remains below four minutes. This modest increase in runtime represents a reasonable trade-off for the substantial reduction in singleton formation achieved with higher branching factors. An additional advantage is the concurrent decrease in peak memory usage as the branching factor increases, improving the accessibility of the method for less modern or resource-limited workstations. Other fingerprint types show same trends (Figures S7-S10). Based on these observations, we proceed with using a branching factor of 1024 and thresholds corresponding to three to four standard deviations above the mean similarity.

**Figure 4.**
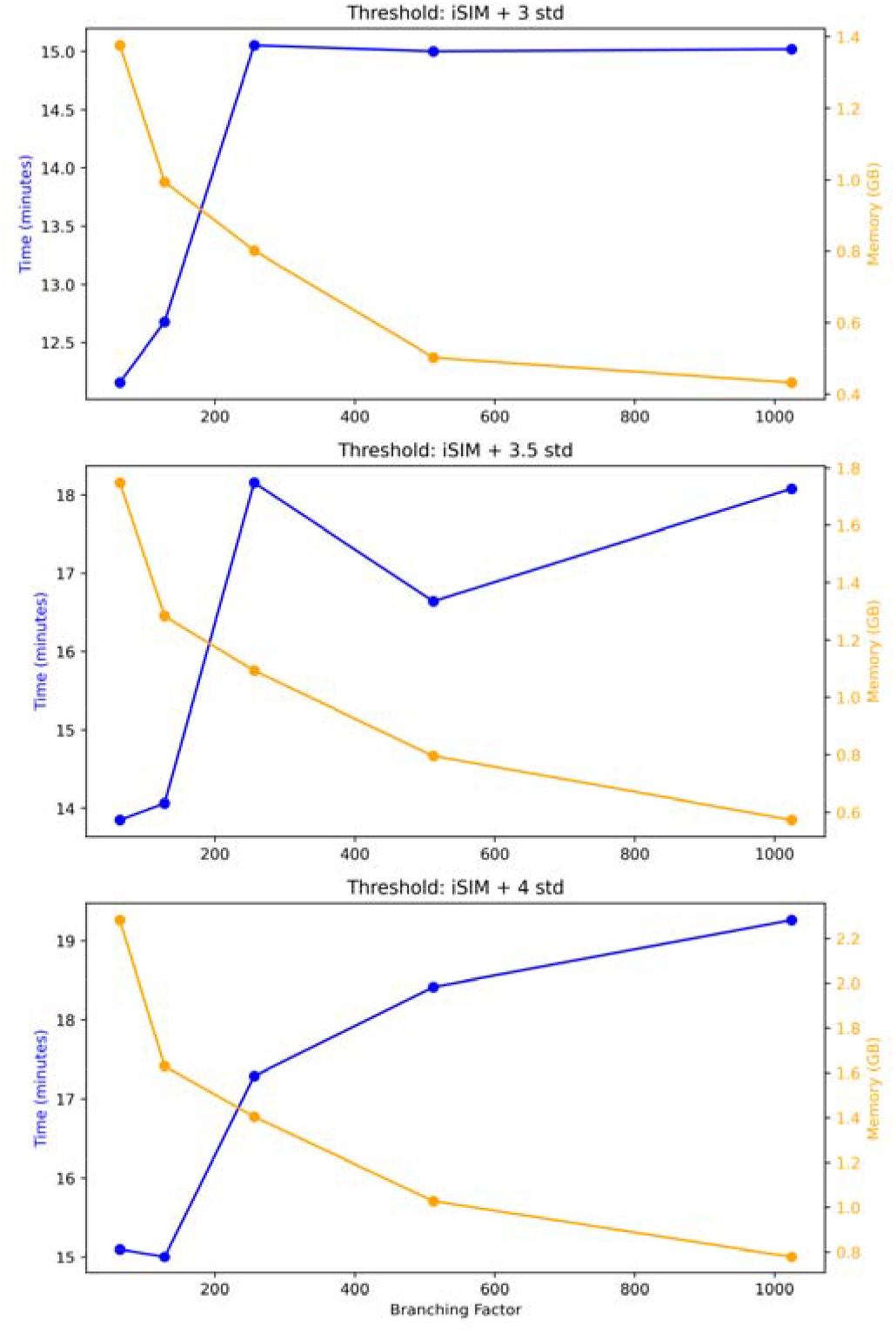
Effect of branching factor and threshold on computing time and memory usage for BitBIRCH clustering of the ChEMBL34 library using ECFP4 2048-bit fingerprints and diameter merging criterion.

We now proceed with a more chemically oriented analysis to evaluate whether the resulting clusters exhibit meaningful structural coherence. In **Table 1**, we report the Maximum Common Substructures (MCS)^20^ for the five most populated clusters, as these regions typically pose the greatest challenge for clustering algorithms and therefore provide a benchmark for assessment. Notably, more complex and chemically informative substructures are recovered when using thresholds of 3.5 and 4 standard deviations above the global iSIM, supporting the selection of these values as the most appropriate. Similar trends are observed for the remaining fingerprint types (See Tables S1-S4). It is worth noting that RDKit fingerprints yield the largest MCS fragments across all representations; however, they are also associated with the highest number of singletons, indicating a trade-off between structural granularity and cluster compactness.

**Table 1.**
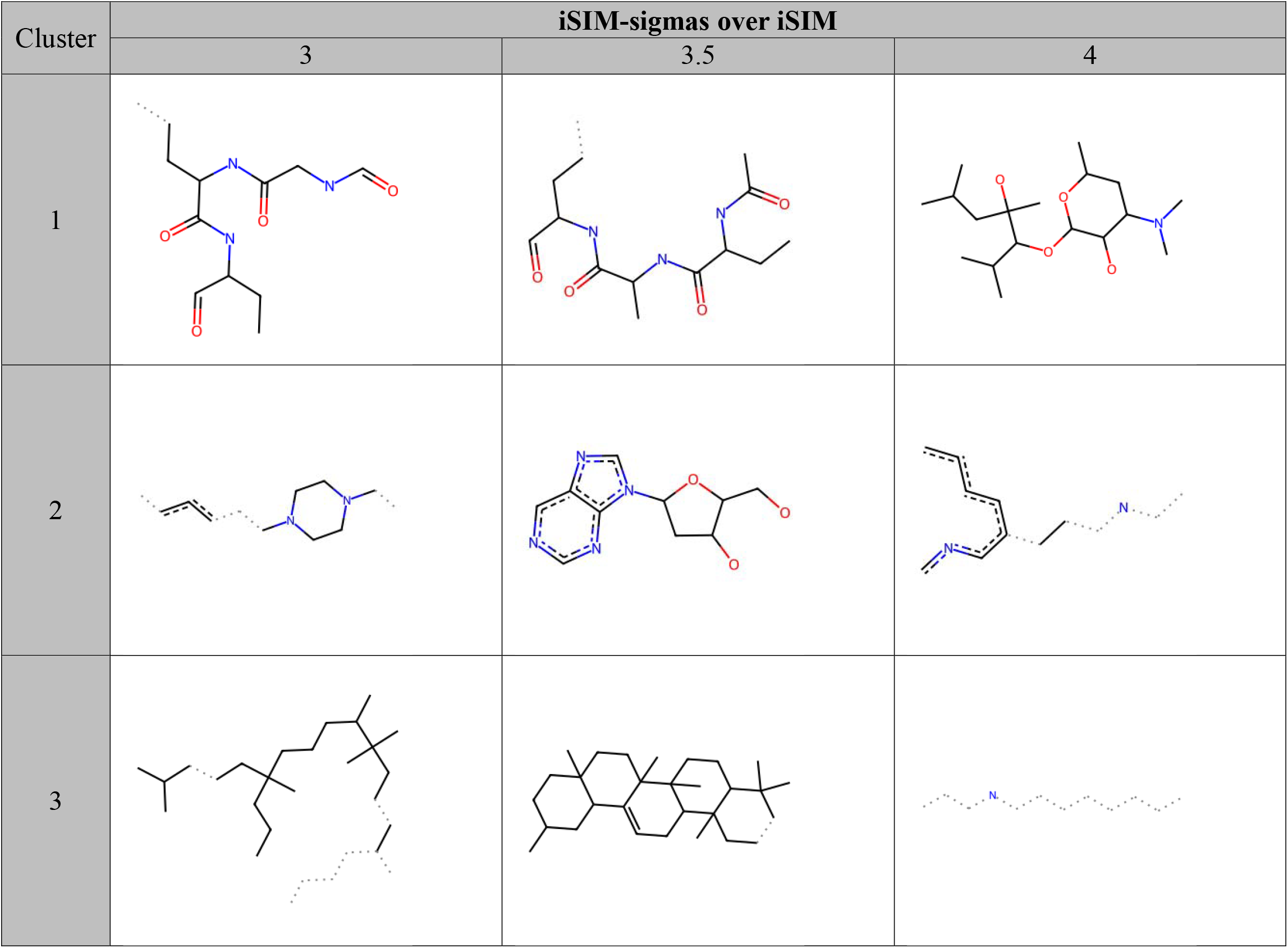
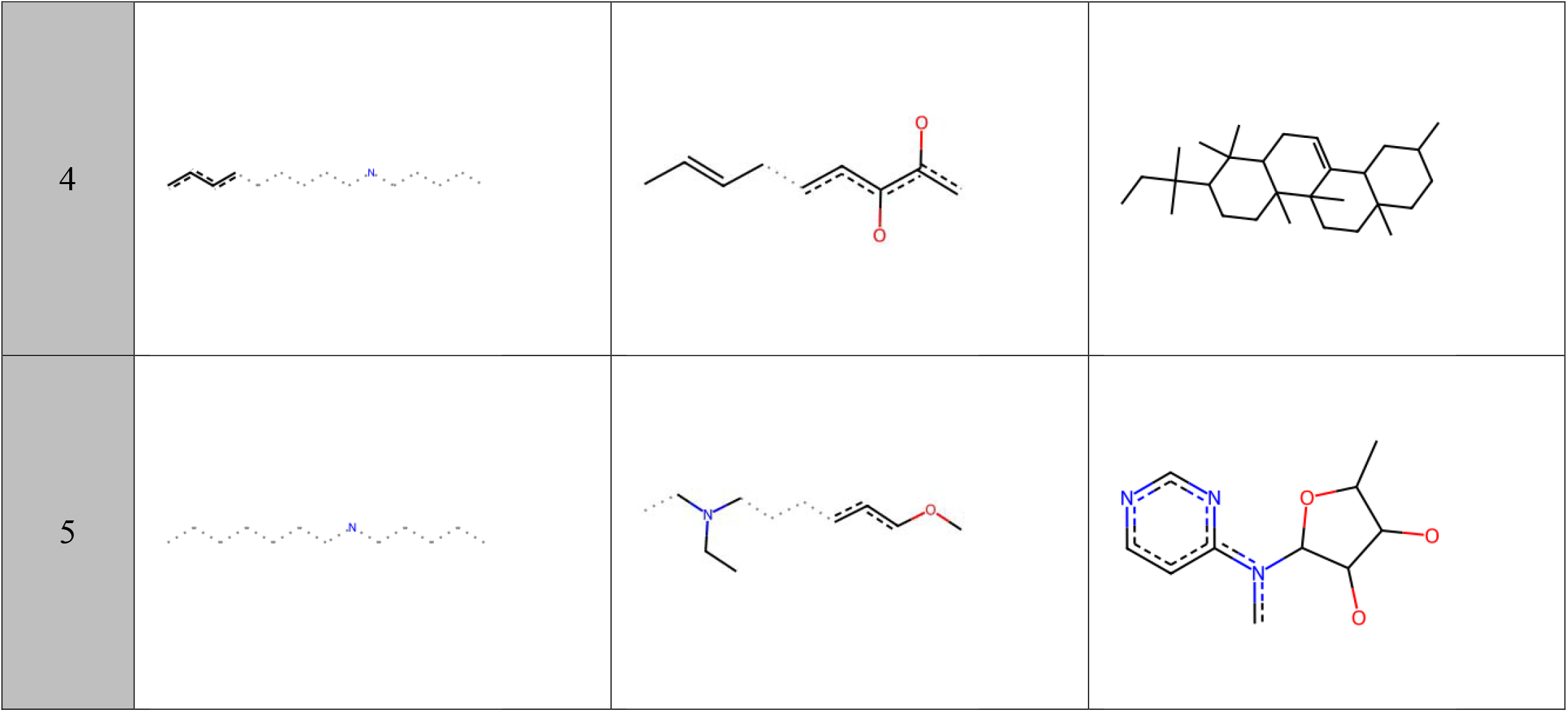
Maximum Common Substructures (MCS) for the five most populated clusters obtained at thresholds of 3, 3.5, and 4 standard deviations above the global iSIM clustered with BitBIRCH with branching_factor = 1024 and diameter merging criterion.

We further observing the structures of the most populated cluster at thresholds of 3 and 3.5 standard deviations, we found mainly peptides and cyclic peptides that not necessarily are similar. (See Figure S11-S13) Given the repetitive nature of amino acid residues, binary fingerprints appear to inadequately capture subtle structural variations within these compounds^21^, leading to artificially cohesive clusters. Excluding peptides from datasets may improve overall clustering performance. **Figure 5** presents the population distributions for the three tested thresholds, showing the absence of a single dominant cluster and a balanced unique scaffold distribution. Nonetheless, the proportion of singletons and small clusters remains relatively high. While the final choice of threshold may depend on the intended application, we recommend 3.5 standard deviations above the mean iSIM as a balanced compromise, as it yields chemically meaningful MCSs and well-populated clusters.

**Figure 5.**
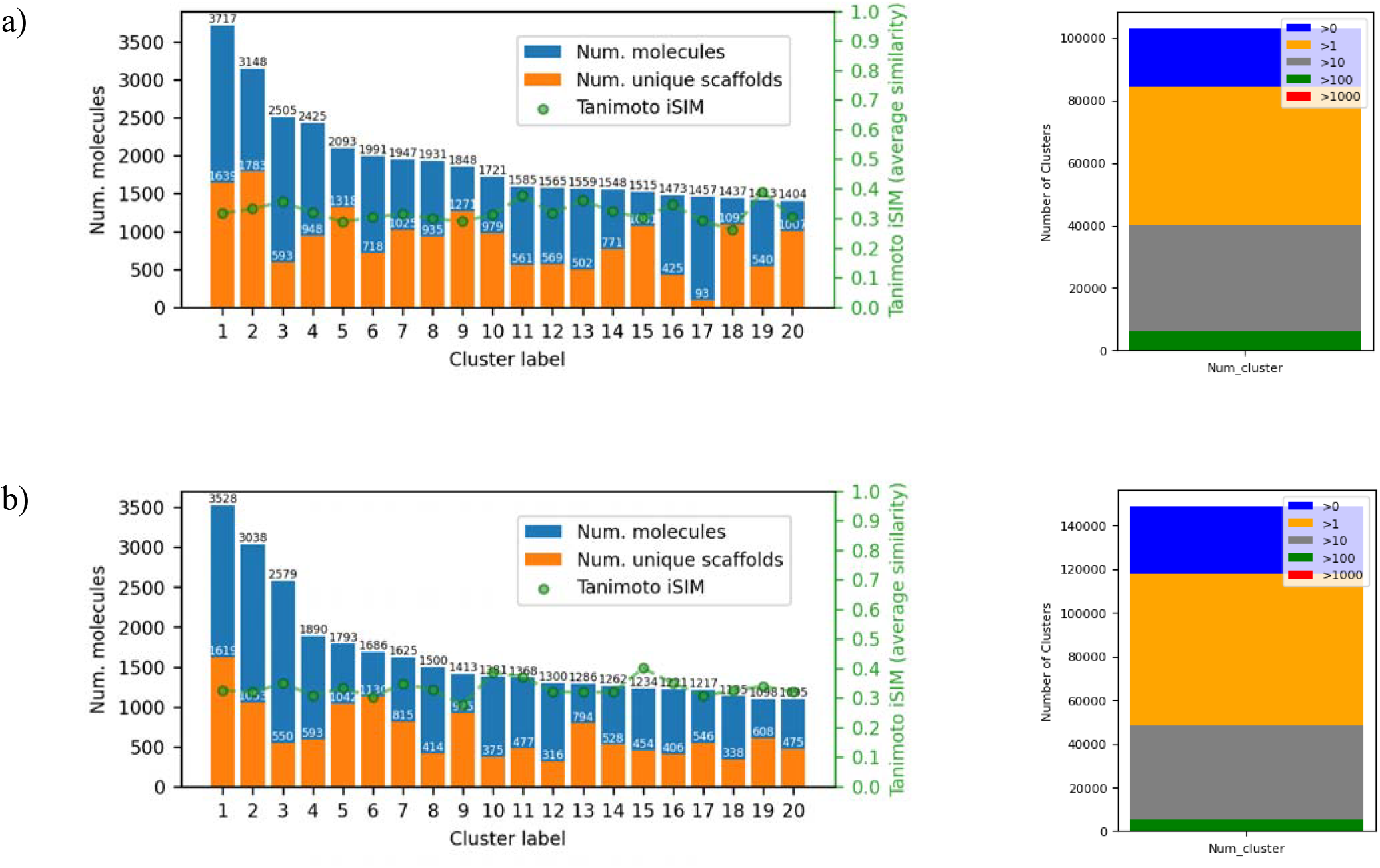
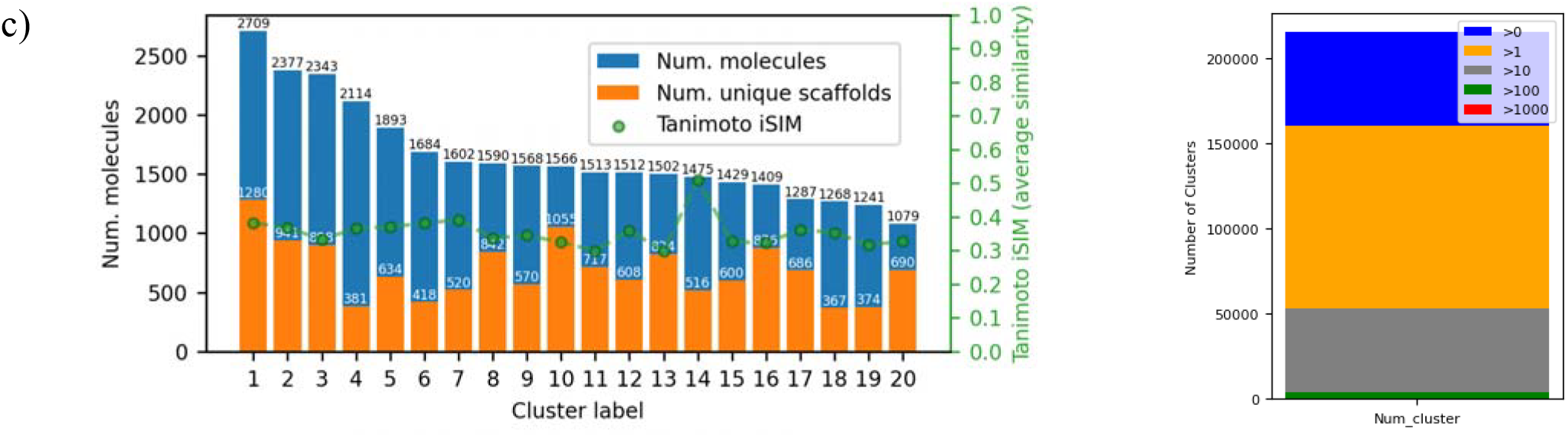
Population and unique scaffold distribution for the top 20 largest clusters and overall cluster population count for the ChEMBL34 library using BitBIRCH (branching factor of 1024) and thresholds of a) 3 b) 3.5 c) 4 standard deviations over iSIM.

As previously discussed, the tree structure of BitBIRCH implies that some molecules or subclusters will not encounter sufficiently similar counterparts during traversal. While increasing the branching factor could alleviate this issue, doing so indiscriminately would significantly increase computational complexity^22^; in the extreme case where the branching factor approaches the number of molecules, the algorithm would degrade to *O(N^2^)*, defeating its primary purpose of efficient scaling.

It the user needs further reduction in certain scenarios, particularly for fingerprints such as RDKit and Atom Pair, where the singleton-to-cluster ratio is not sufficiently diminished by increasing the branching factor alone. (See Figure S3-S4) To address this limitation, we implemented a re-clustering strategy that exploits the flexibility of the BitBIRCH framework. This approach, enabled via the newly introduced `recluster_inplace` method, re-clusters the partitions generated in the initial step, thereby increasing the probability of merging structurally related molecules that were previously separated due to the constraints imposed by the tree structure.

During this re-clustering process, an additional threshold constraint can be applied to prevent the formation of overly large or diffuse clusters, thus preserving the quality of the original partition. The procedure can be repeated iteratively until no further changes in cluster assignments are observed. Importantly, each re-clustering step is computationally less demanding than the initial clustering, as it operates on BitFeatures derived from the previous pass rather than on the full set of molecular fingerprints.

In **Figure 6**, we illustrate the evolution of cluster population, number of singletons, and singleton-to-cluster ratio across successive re-clustering iterations of the initial ChEMBL clustering. At each iteration, we evaluated two scenarios: adding one standard deviation to the threshold and applying no additional threshold. In both cases, the number of clusters and singletons begins to stabilize around the fifth iteration, indicating convergence of the clustering process. Similar trends were observed across other fingerprint types.

**Figure 6.**
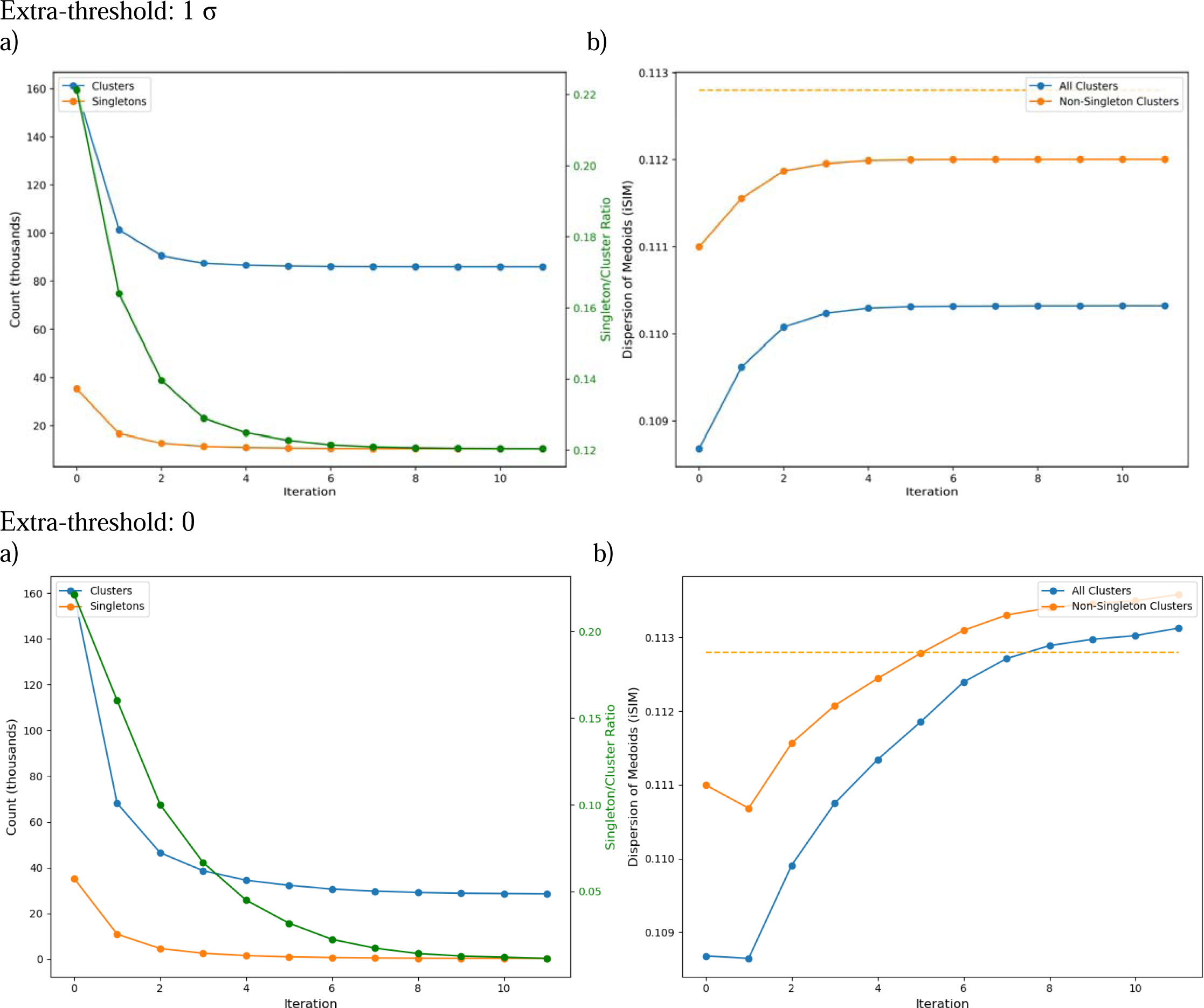
Evolution of (a) the number of clusters, number of singletons, and singleton-to-cluster ratio, and (b) the iSIM of the cluster medoids as a function of successive re-clustering iterations. Results correspond to the ChEMBL34 library represented with ECFP4 2048-bit fingerprints and clustered using BitBIRCH with a branching factor of 1024 and a threshold set to 3.5 standard deviations above the global iSIM of the dataset with extra-threshold of one standard deviation and zero.

Notably, the behavior of the iSIM of the medoids differs depending on whether an extra threshold is applied. When one standard deviation is added, the medoid iSIM reaches a plateau early in the process, whereas in the absence of an extra threshold it continues to increase across iterations. In the extra-threshold scenario, the medoid iSIM remains below the global iSIM of the full dataset, approaching it closely (within the third decimal place), while without the additional threshold this global value is surpassed. Introducing an extra threshold therefore appears beneficial for mitigating over merging, as it prevents the formation of clusters with excessively similar medoids. However, with no extra-threshold the number of clusters and singletons is significantly reduced.

A closer analysis of the cluster population comparing the effect of the no and with extra-threshold in the re-clustering steps is illustrated in Figure 7. Here we can see that the populations of the clusters are much larger when no extra-threshold is added; in consequence we can see that the number of singletons and small clusters is significantly reduced. We also want to point out that the proportion of unique scaffolds to the cardinality of the clusters maintains a similar trend in both instances, suggesting that there is not much detriment to the clustering in terms of chemical structures. In Table S5, we calculated the MCS for the top 20 clusters, we can see that for both instances we obtain similar core MCS even with such difference in the extra-threshold, but such a big difference in populations. Taking a closer look at some clusters in Tables S6-S7, we see that where the difference lays is in the similarity of the decorations of the core MCS structures; with no extra-threshold clusters with the same MCS are merged, however there is a high diversity of decorations. In Tables S8-S9 and Figure S14 we can see some examples of when the re-clustering with extra-threshold can effectively merge singletons and small clusters that are structurally similar. With an extra-threshold the substituents attached to the MCS are more similar as it is illustrated in each of the subclusters in Tables S6-S7. Additionally, because of the size a few random molecules with no chemical significance can be included if no extra-threshold is used.

**Figure 7.**
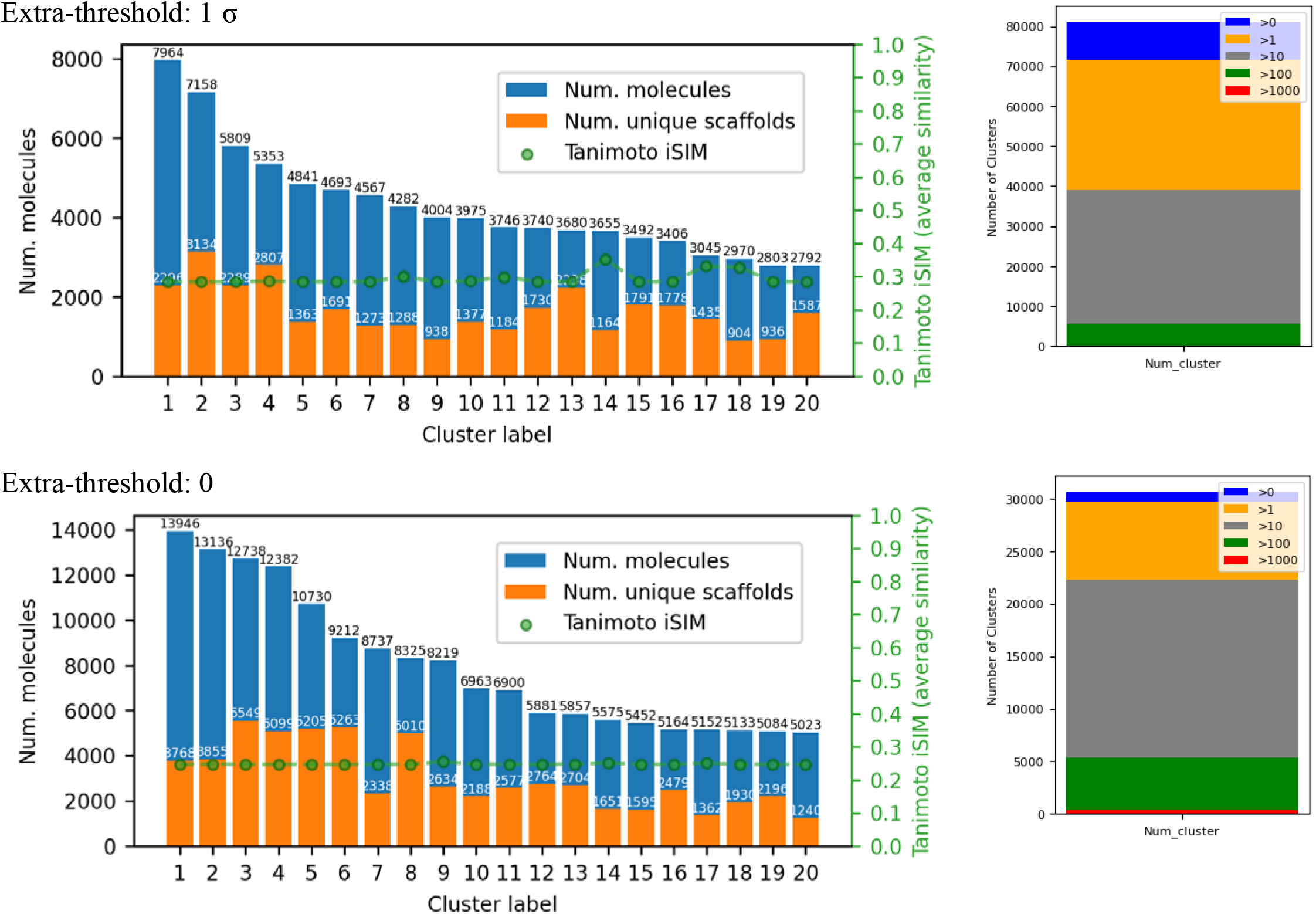
Population and unique scaffold distribution for the top 20 largest clusters and overall cluster population count for the ChEMBL34 library using BitBIRCH (branching factor of 1024, threshold of 3.5 standard deviations over iSIM) and five re-clustering iterations.

The use of an additional threshold during re-clustering should be guided by the specific objectives of the usage of BitBIRCH. If the primary goal is the identification of molecules sharing a common core, the application of an extra threshold may be unnecessary. In contrast, for tasks that require more homogeneous clusters, such as QSAR studies, the use of a higher threshold combined with an additional extra-threshold becomes advantageous to ensure tighter clusters.

Although the strategies described above substantially improve clustering quality across most tested fingerprint types, RDKit fingerprints represent a notable exception. For this representation, a large, loosely defined cluster persists, and the number of singletons remains higher compared to other fingerprints. While post-processing and refinement strategies can further mitigate these effects, some clustering patterns are intrinsically linked to the molecular encoding itself. As previously discussed, RDKit fingerprints tend to generate clusters characterized by larger and more structurally detailed MCS, which rationalizes the continued presence of numerous small clusters even after applying the proposed methodology.

Consequently, RDKit fingerprints may be particularly valuable for studies requiring highly detailed and fine-grained molecular clustering. Strategies for partitioning the large cluster have been explored in previous work^8^; we recommend that, for RDKit fingerprints, the fragmentation and redistribution of this dominant cluster be performed prior to the re-clustering step. Additional details are provided in the specific notebook for RDKit fingerprints.

This strategy was further validated on two additional databases, for which the observed trends agreed with the results obtained for the ChEMBL34 case study. Details are included in the Supporting Information.

## Conclusions

Clustering threshold selection cannot be universally prescribed, as it depends on both the molecular representation, the dataset, and objectives of the user. In this communication, we establish a systematic, data-driven framework for the rational parameterization of BitBIRCH. Our results demonstrate that coherent clustering behavior and effective reduction are achieved when the threshold is set between three and four standard deviations above the average similarity of the dataset. The combined use of iSIM and iSIM-σ provides an efficient, linear scaling and accurate method for calculating the optimal threshold in each instance.

We further show that increasing the branching factor significantly decreases singleton formation, reinforcing its critical role in controlling number of clusters and singletons. We recommend employing branching factors as high as computationally feasible without sacrificing efficiency; for the library studied here, a value of 1024 provides an optimal balance between performance and clustering quality, although this parameter should be scaled according to dataset size.

In scenarios where the user needs to address the persistence of small clusters and singletons, we propose a re-clustering strategy that promotes further consolidation of structurally related molecules. Up to five re-clustering iterations are recommended, as beyond this point no substantial changes in cluster composition are observed. The use of an additional similarity threshold during this process should be dictated by the intended application: while omitting the extra-threshold facilitates the identification of broader structural relationships, incorporating a value equivalent to one standard deviation helps prevent over merging and preserves more homogeneous clusters.

Overall, this work provides a practical and statistically founded set of guidelines for the application of BitBIRCH, balancing clustering efficiency, structural coherence, and computational performance. While the proposed parameters reflect optimal settings for common clustering objectives, the framework presented here gives users a better idea on how to tailor the method to alternative goals.

## Methods

Jupyter notebooks with workflow to obtain the analysis in this paper are included in our GitHub repository https://github.com/mqcomplab/bblean. Binary fingerprints were generated using RDKit^15^. MCS were determined by a random down-sample of each cluster with a threshold of 0.75 using RDKit’s implementation. ChEMBL^16^ library was obtained from publicly available download tool https://www.ebi.ac.uk/chembl/explore/compounds/, compounds flagged as inorganic were discarded from this study. ‘diameter’ merging criterion was used for all the BitBIRCH clustering in this work.

## ACKNOWLEDGEMENTS

We thank support from the National Institute of General Medical Sciences of the National Institutes of Health under award number R35GM150620.

