## Supplementary material for "Best practices to cluster large molecular libraries": Supplemtary Information

**Section 1:** ChEMBL34 library (excluding entries flagged as inorganic, *n* = 2,764,449)

AP

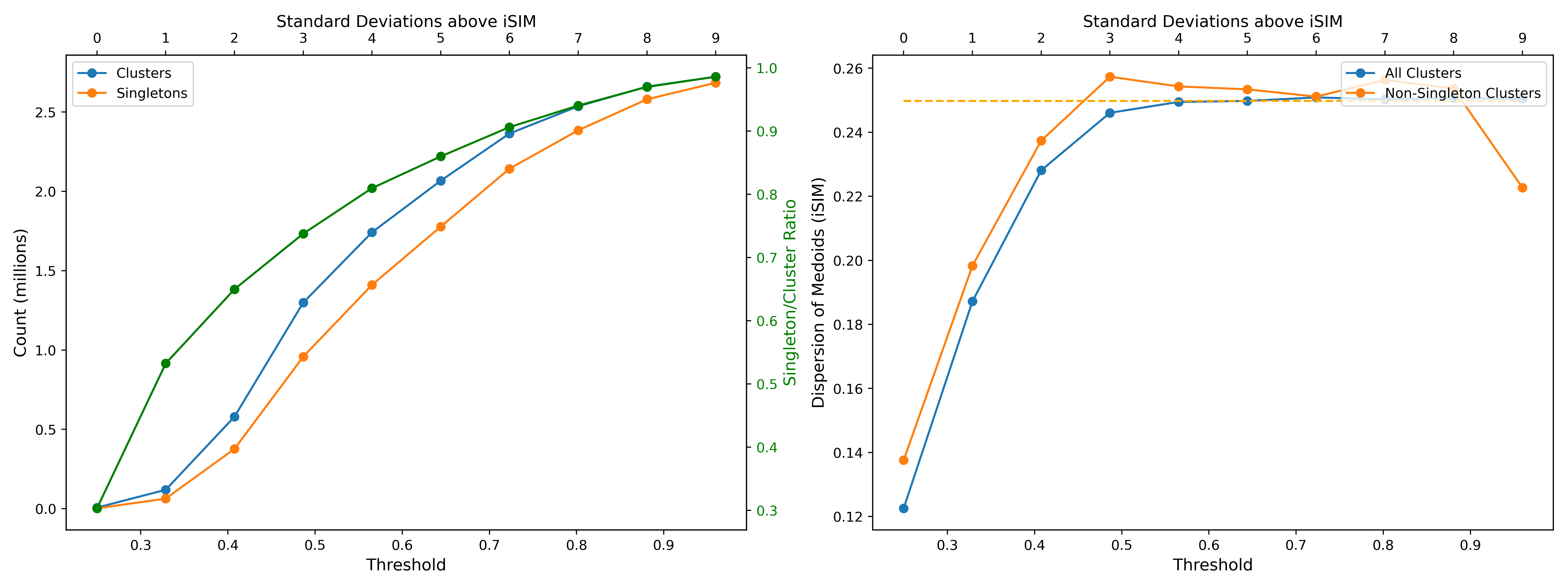

RDKIT

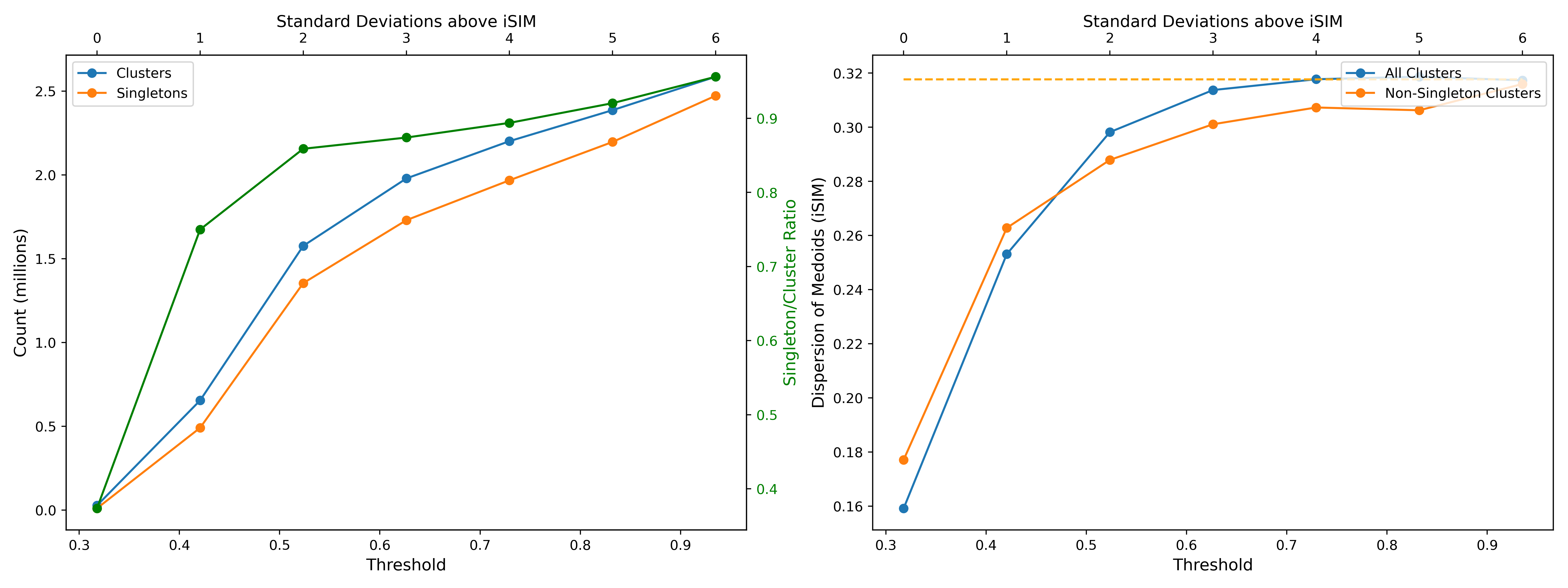

ECFP6

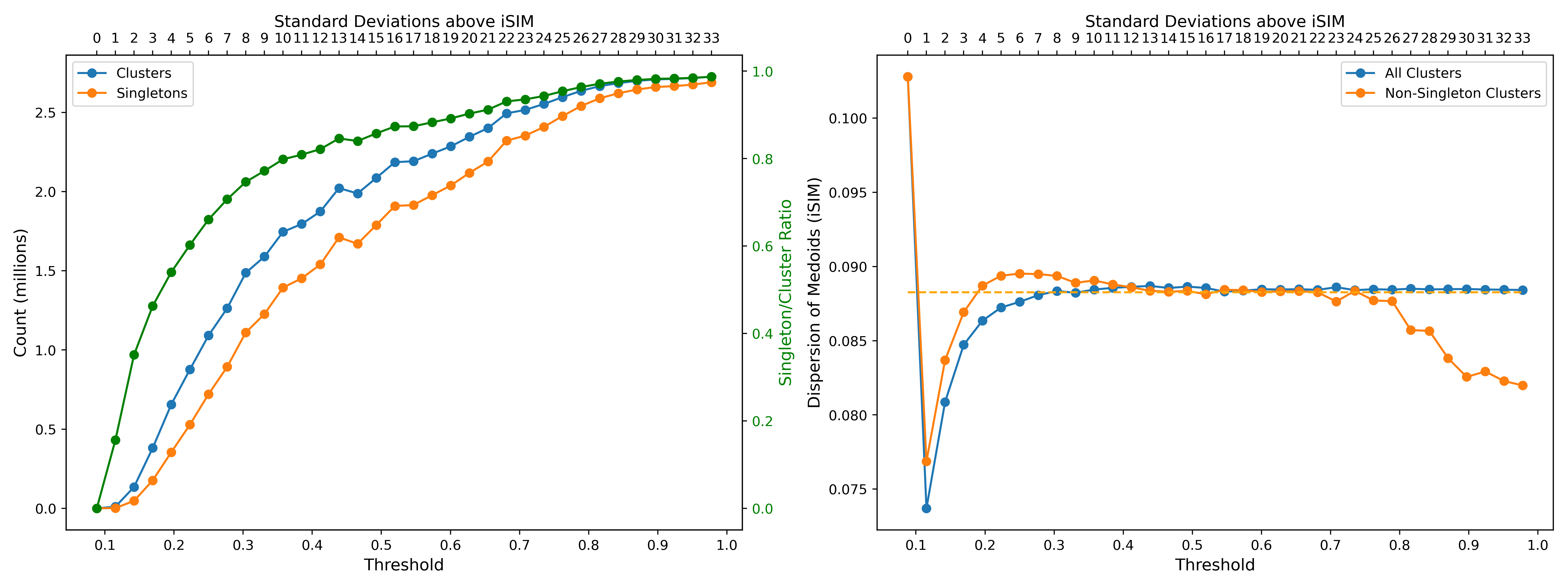

Topological Torsion

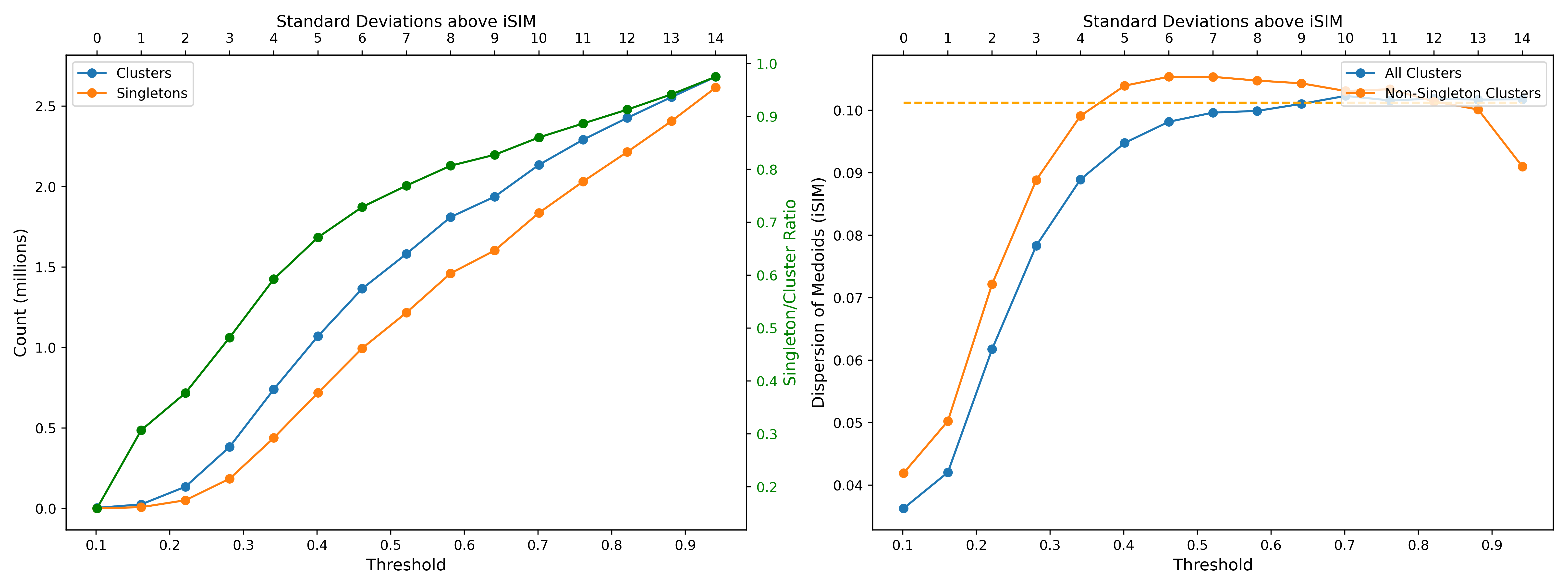

**Figure S1.** Variation of the number of clusters, singletons, singleton-to-cluster ratio; and iSIM of the medoids for the ChEMBL34 library represented with different types of 2048-bit fingerprints. iSIM of the global library is indicated with dashed line.

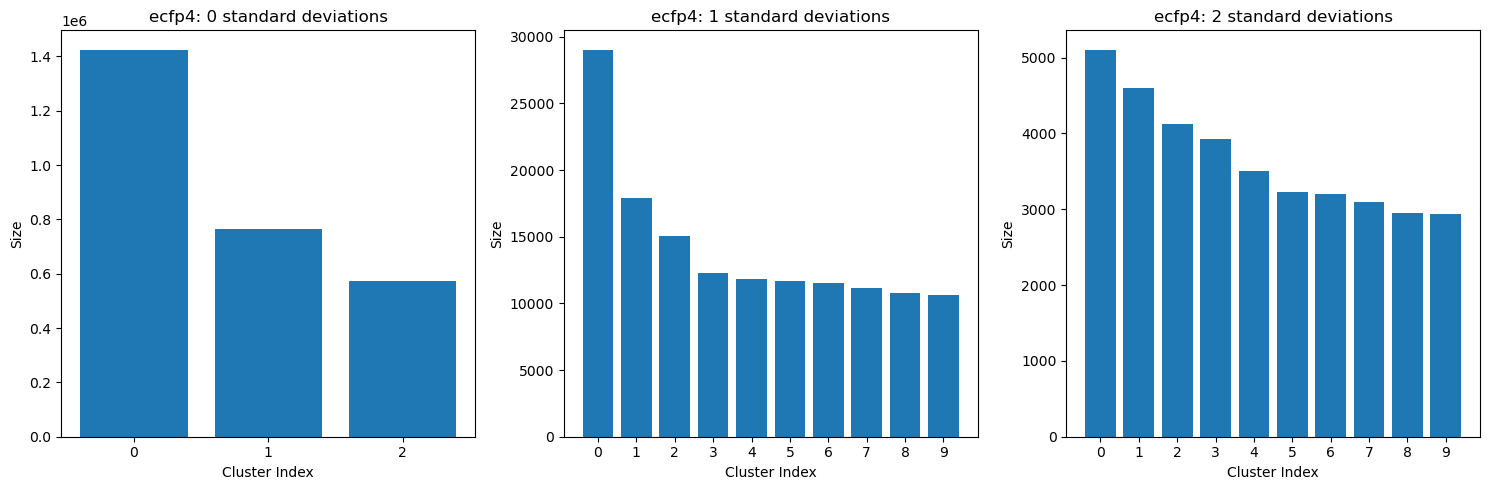

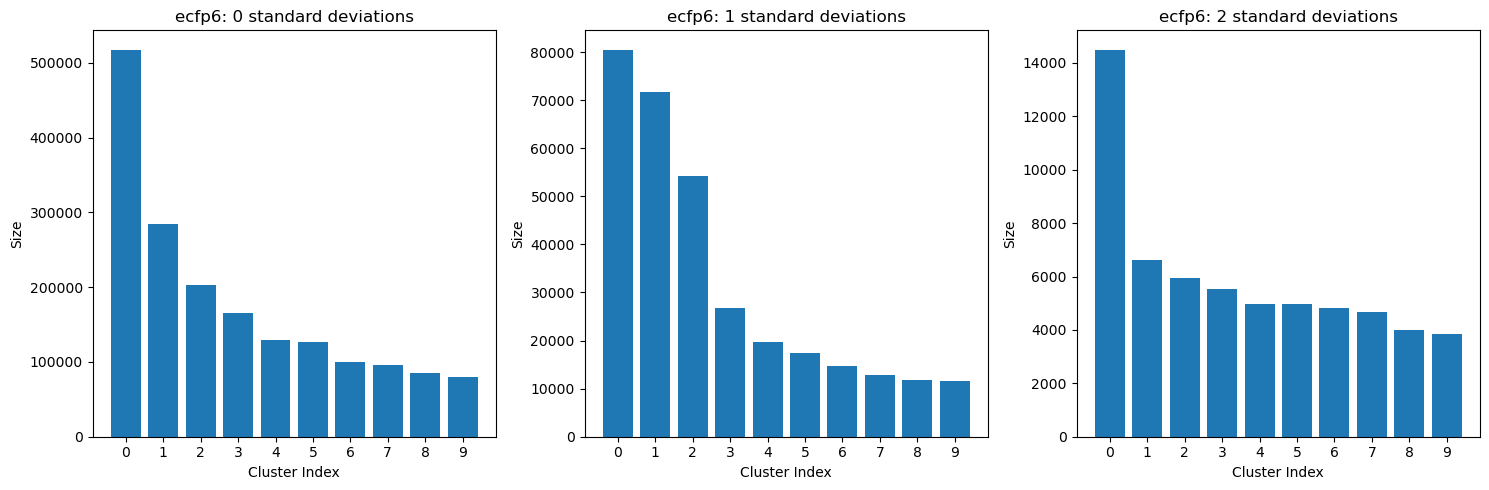

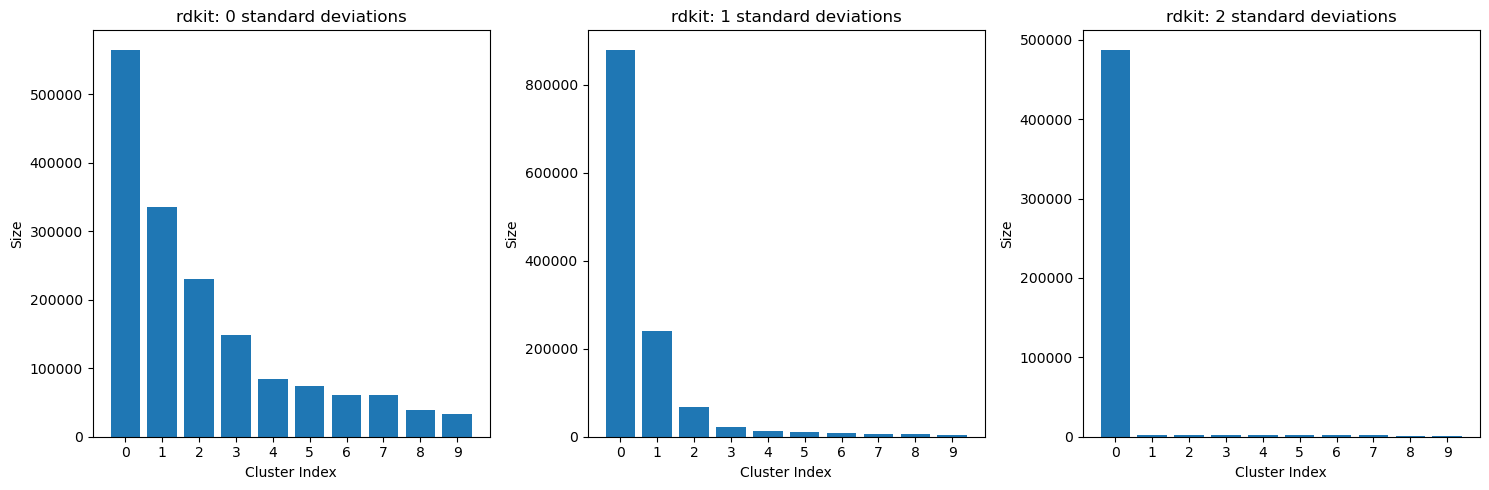

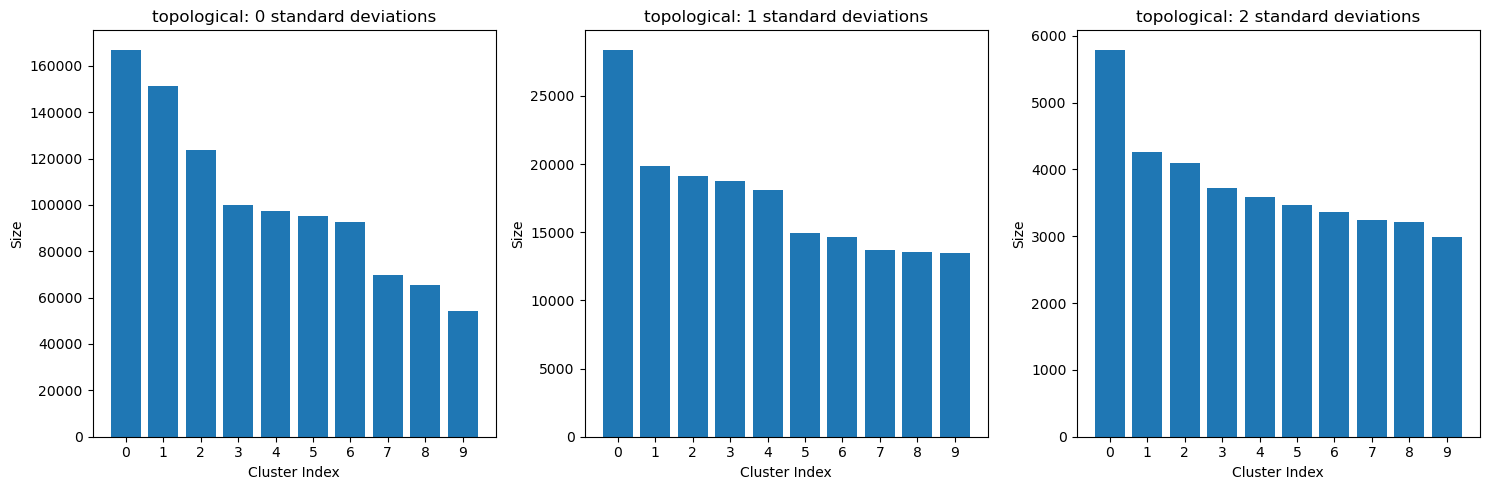

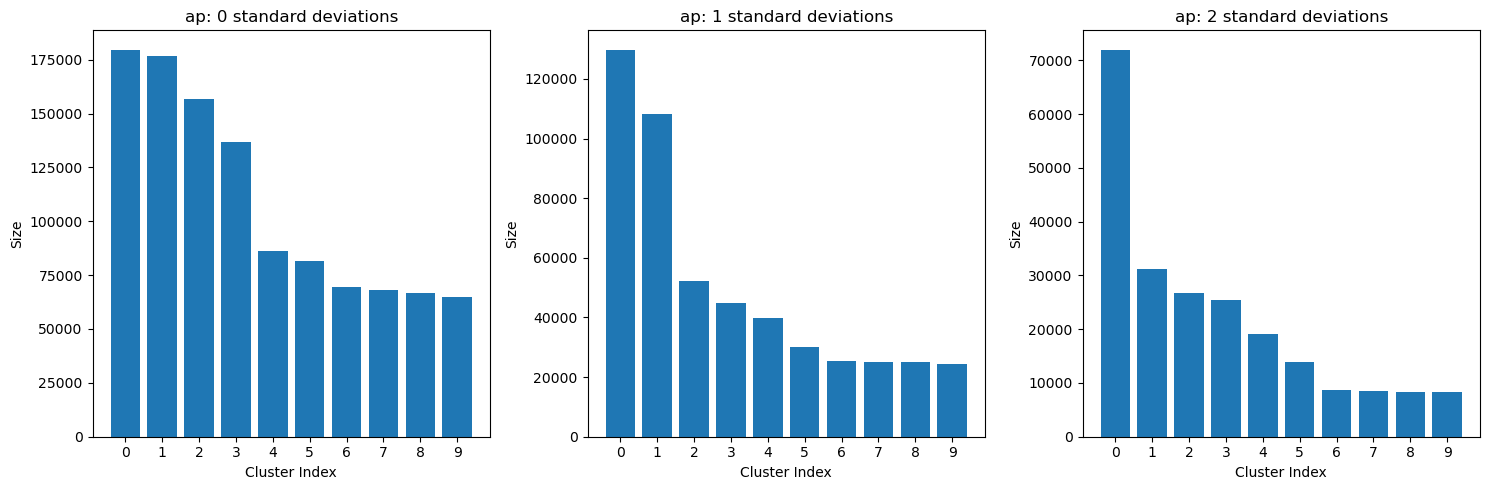

**Figure S2.** Populations of the top 10 most populated clusters for all the tested fingerprints with thresholds of 0, 1 and 2 standard deviations above the global iSIM.

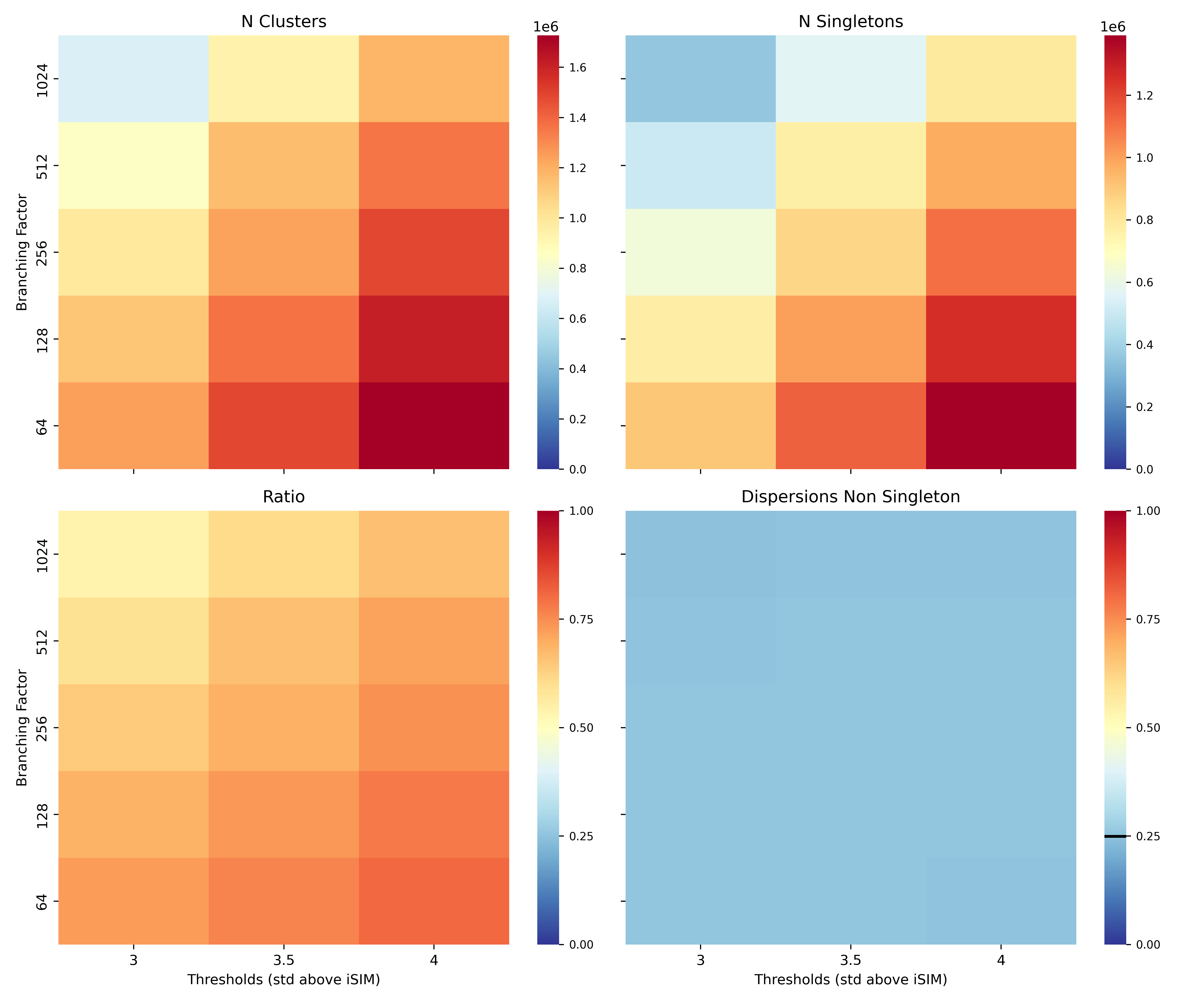

**Figure S3.** Variation of the number of clusters, number of singletons, ratio singleton/clusters and the dispersion (iSIM of medoids) of non-singleton clusters with respect to the branching factor and the thresholds, for the ChEMBL34 library represented with 2048-bit binary AP fingerprints. iSIM of the whole library is marked with a black line in the dispersions panel.

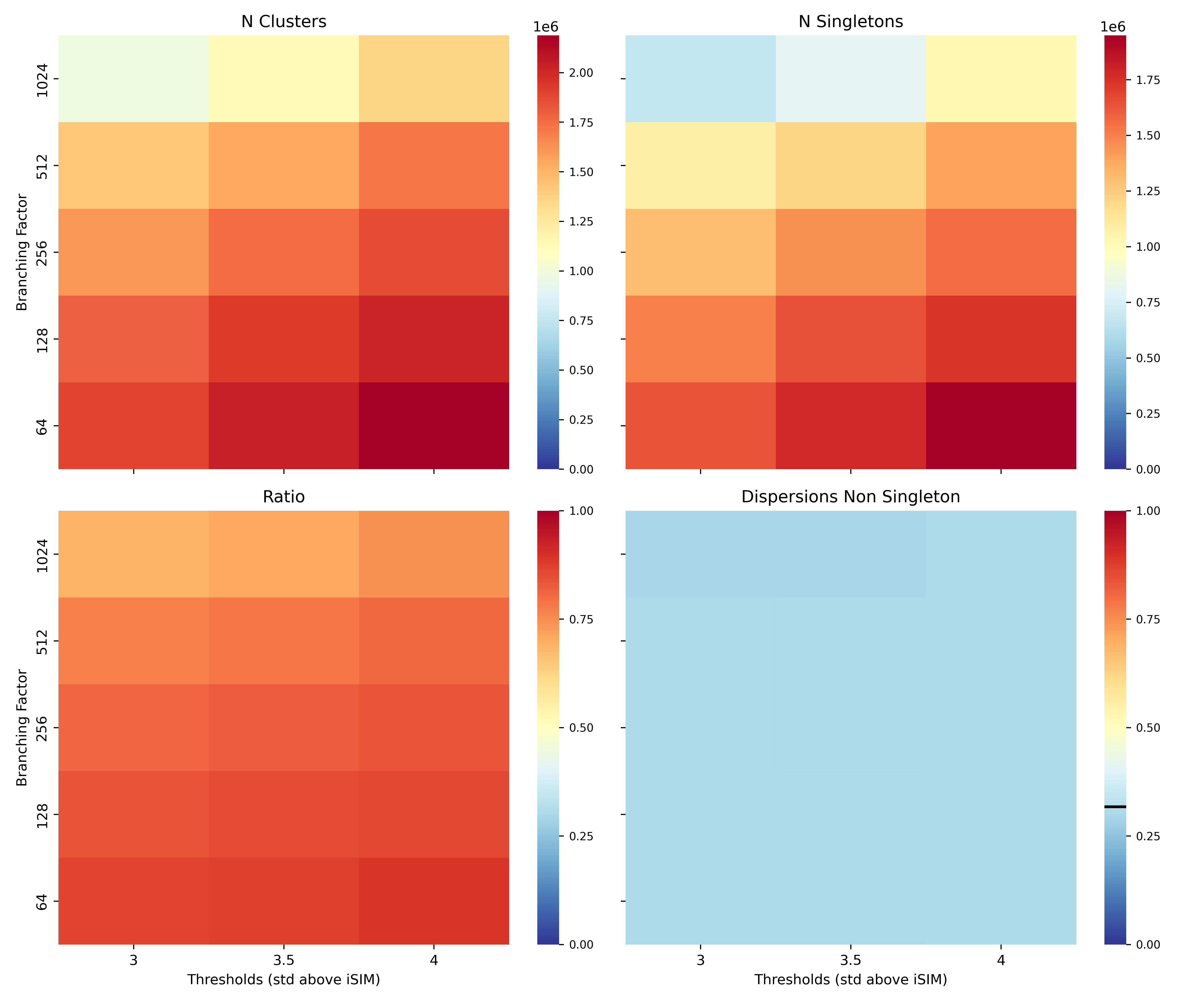

**Figure S4.** Variation of the number of clusters, number of singletons, ratio singleton/clusters and the dispersion (iSIM of medoids) of non-singleton clusters with respect to the branching factor and the thresholds, for the ChEMBL34 library represented with 2048-bit binary RDKIT fingerprints. iSIM of the whole library is marked with a black line in the dispersions panel.

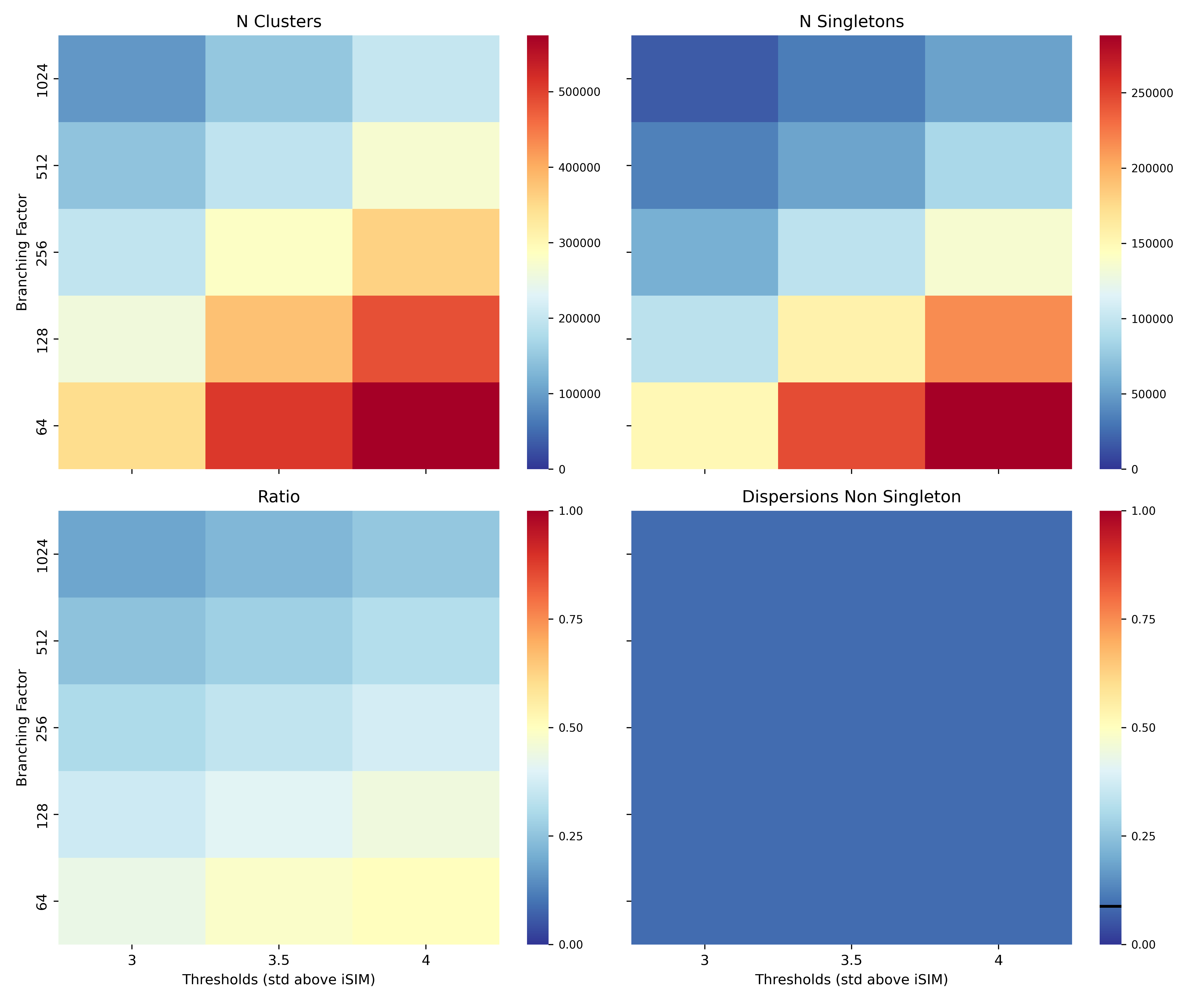

**Figure S5.** Variation of the number of clusters, number of singletons, ratio singleton/clusters and the dispersion (iSIM of medoids) of non-singleton clusters with respect to the branching factor and the thresholds, for the ChEMBL34 library represented with 2048-bit binary ECFP6 fingerprints. iSIM of the whole library is marked with a black line in the dispersions panel.

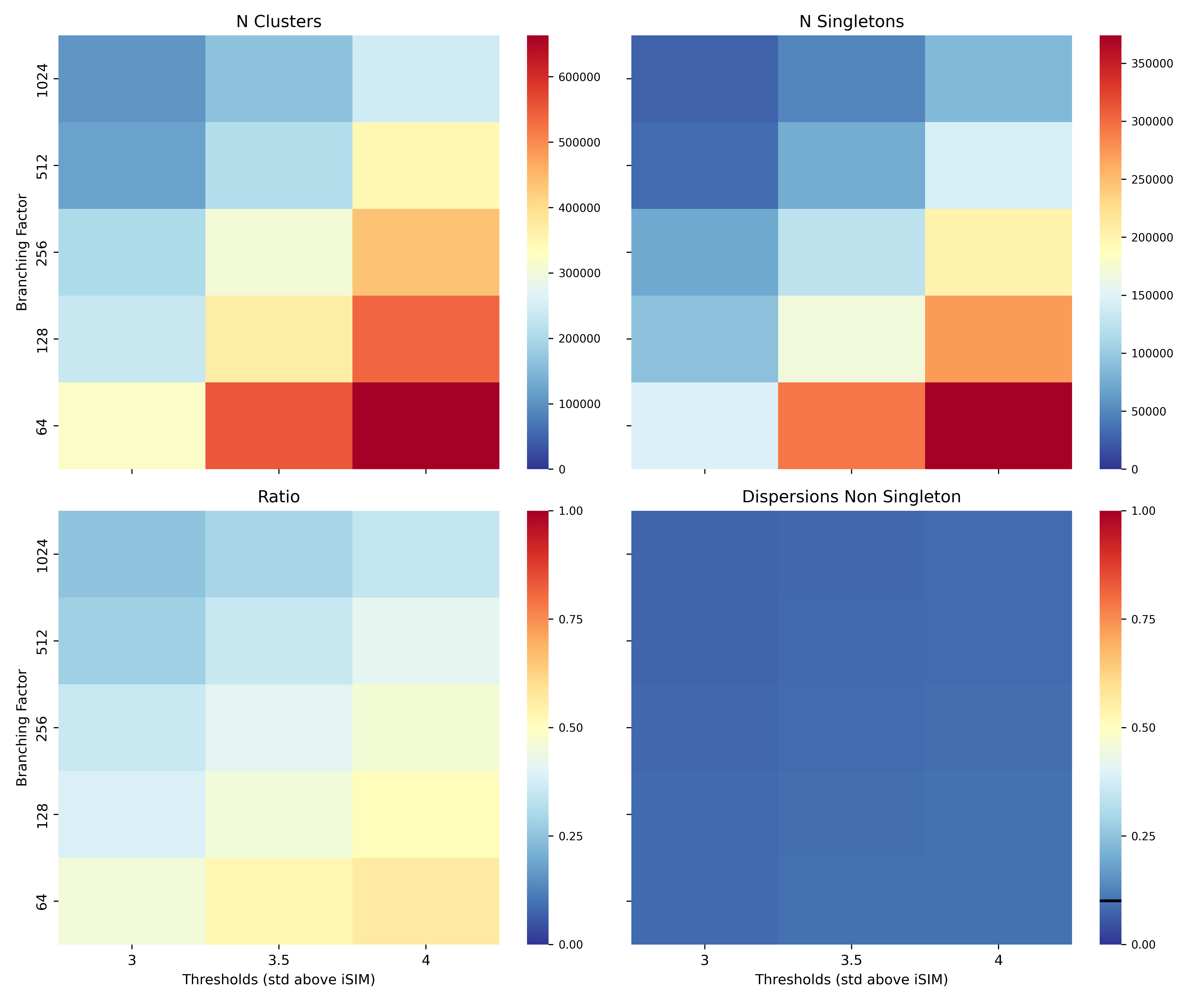

**Figure S6.** Variation of the number of clusters, number of singletons, ratio singleton/clusters and the dispersion (iSIM of medoids) of non-singleton clusters with respect to the branching factor and the thresholds, for the ChEMBL34 library represented with 2048-bit binary Topological Torsion fingerprints. iSIM of the whole library is marked with a black line in the dispersions panel.

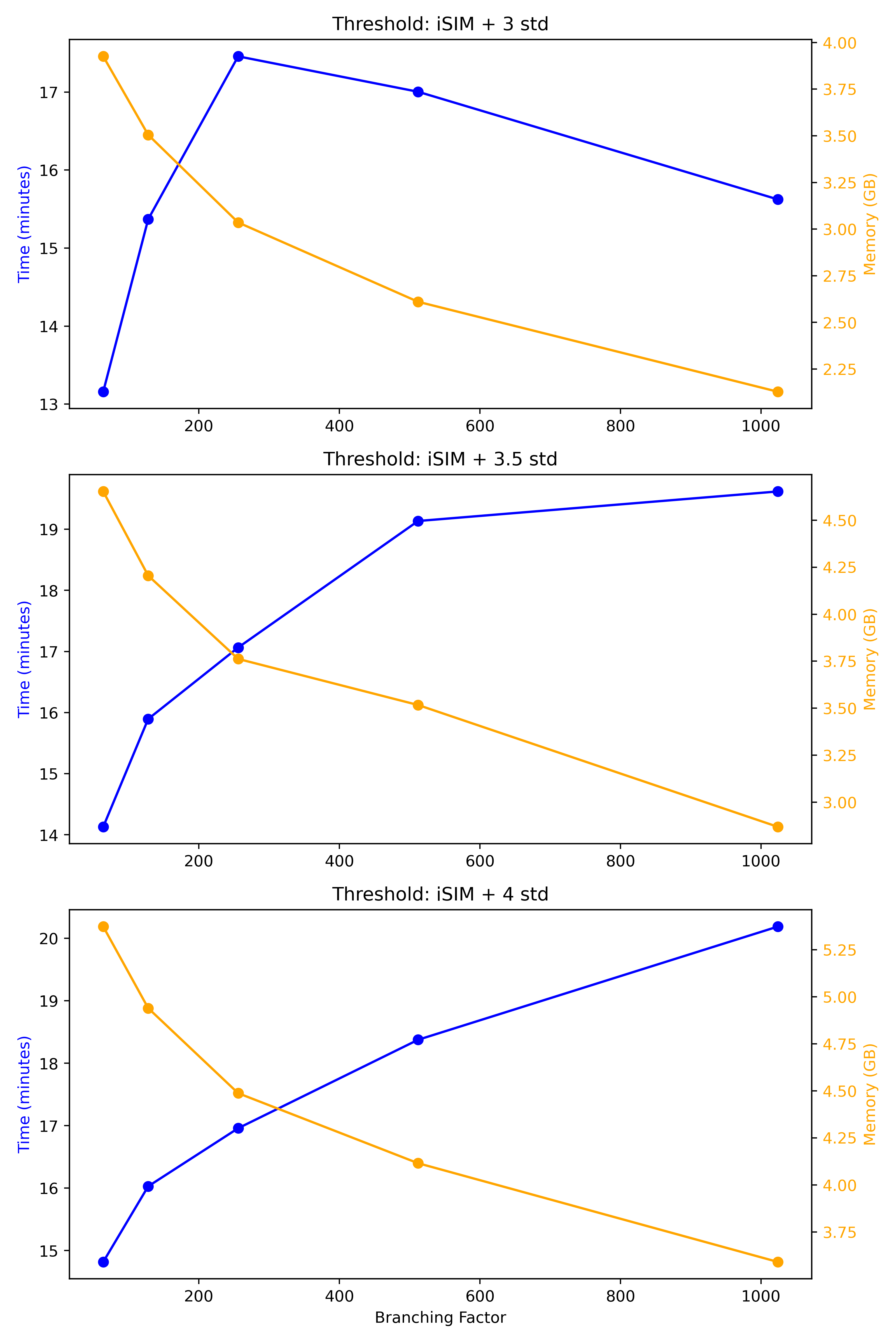

**Figure S7.** Effect of branching factor and threshold on computing time and memory usage for BitBirch clustering of the ChEMBL34 library using AP 2048-bit fingerprints and diameter merging criterion.

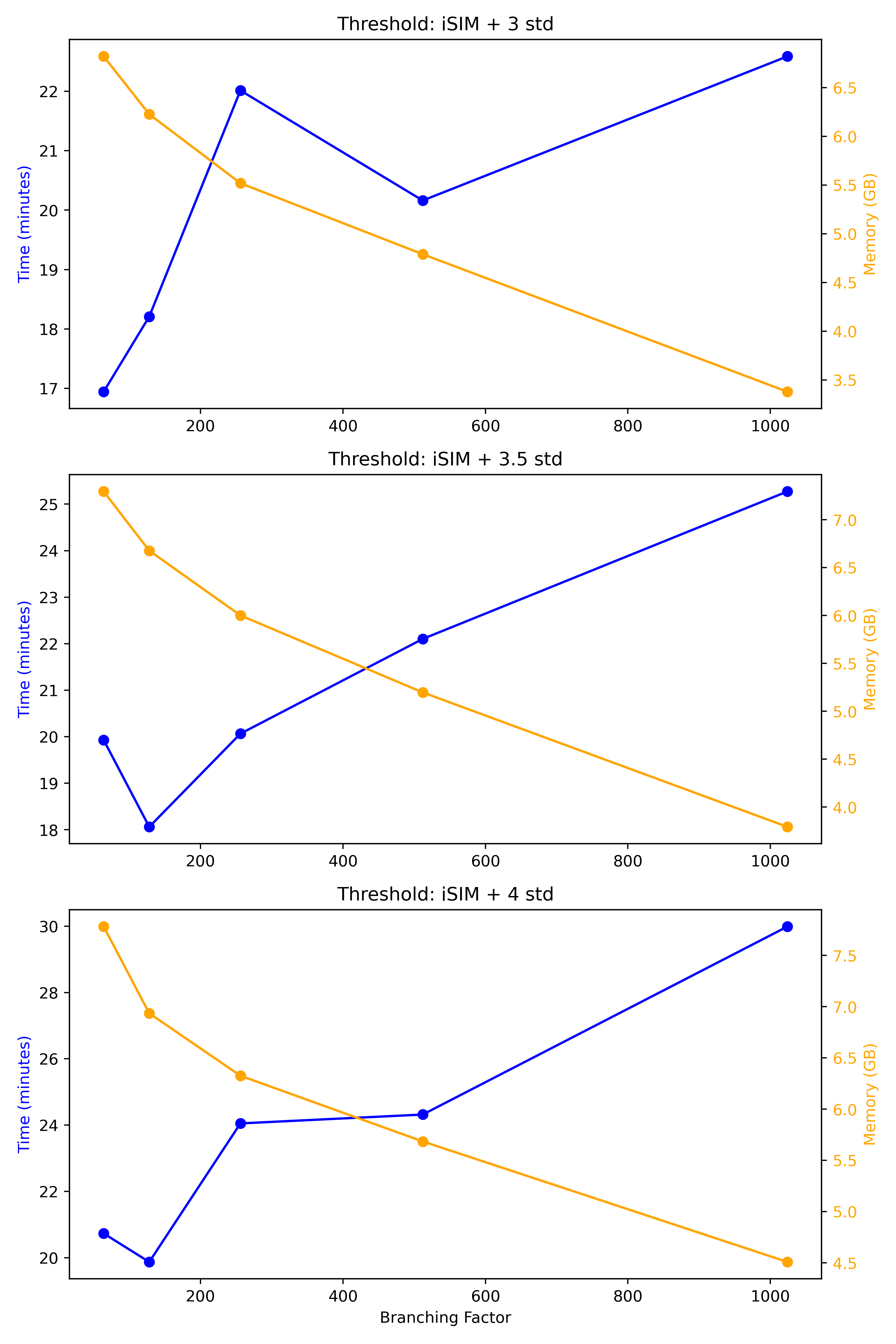

**Figure S8.** Effect of branching factor and threshold on computing time and memory usage for BitBirch clustering of the ChEMBL34 library using RDKIT 2048-bit fingerprints and diameter merging criterion.

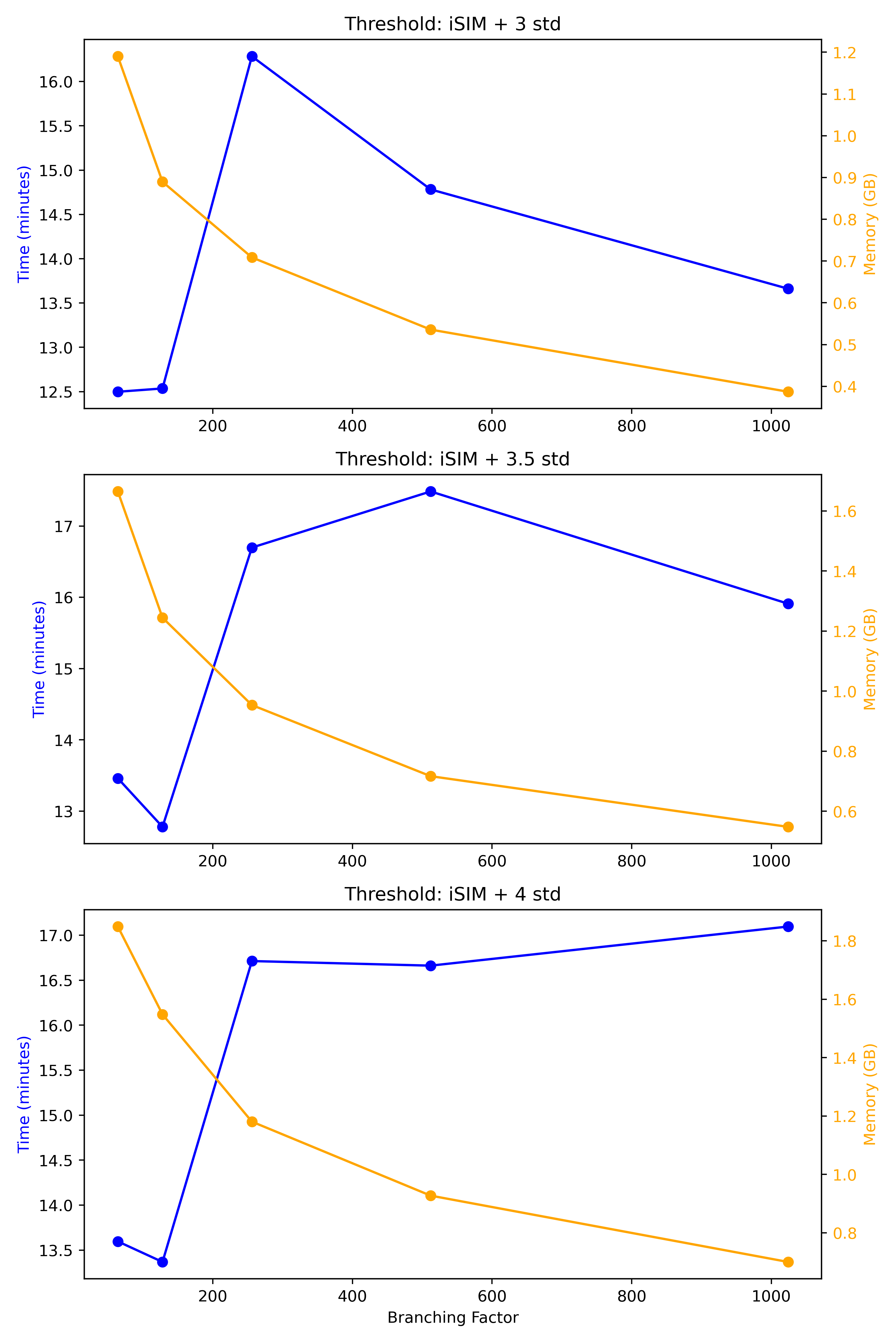

**Figure S9.** Effect of branching factor and threshold on computing time and memory usage for BitBirch clustering of the ChEMBL34 library using ECFP6 2048-bit fingerprints and diameter merging criterion.

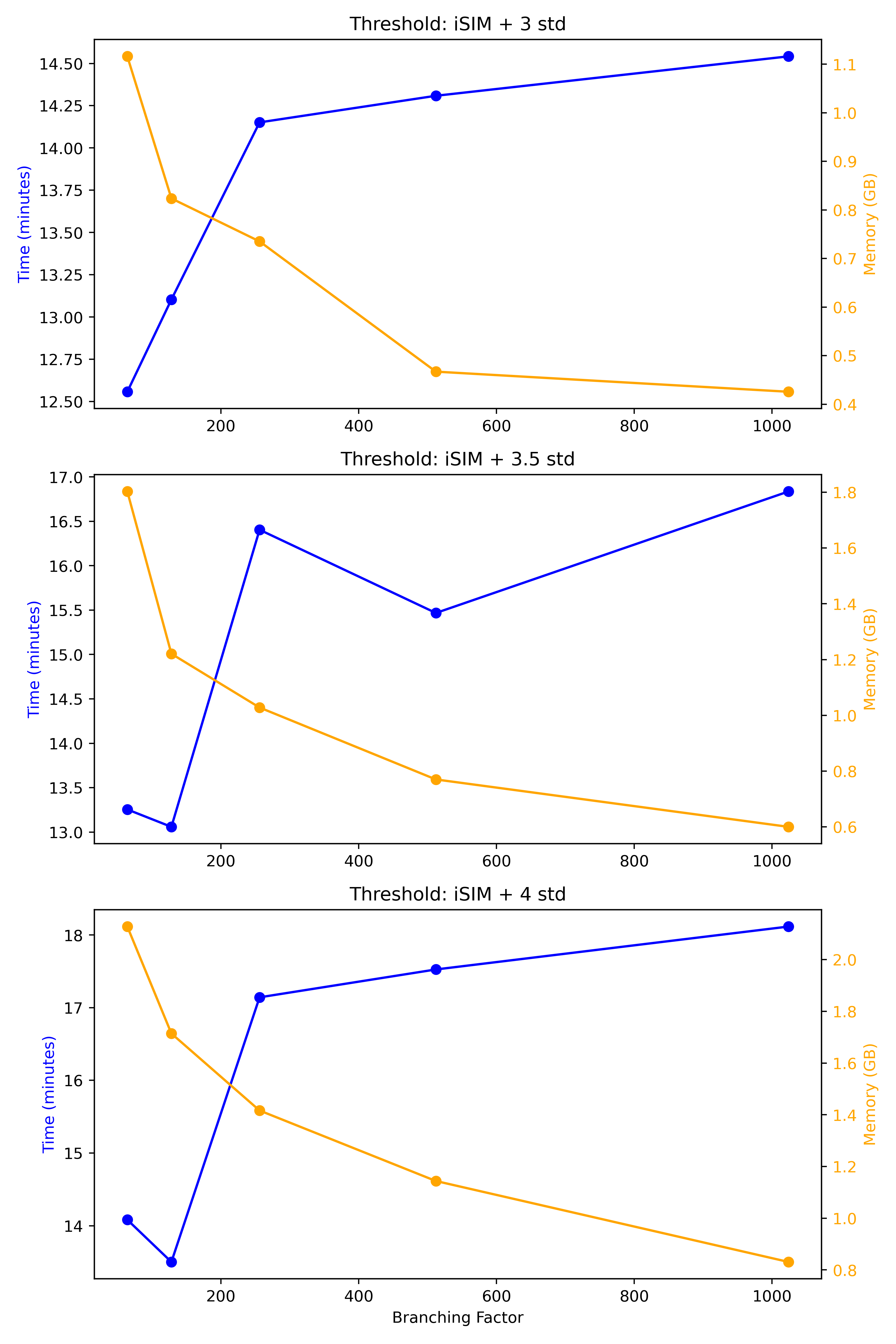

**Figure S10.** Effect of branching factor and threshold on computing time and memory usage for BitBirch clustering of the ChEMBL34 library using Topological Torsion 2048-bit fingerprints and diameter merging criterion.

**Table S1.** Maximum Common Substructures (MCS) for the five most populated clusters obtained at thresholds of 3, 3.5, and 4 standard deviations above the global iSIM clustered with BitBirch with branching_factor = 1024 and diameter merging criterion for ECFP6 fingerprints.

| Cluster | iSIM-sigmas added to iSIM | | |
| --- | --- | --- | --- |
|  | 3 | 3.5 | 4 |
| 1 | 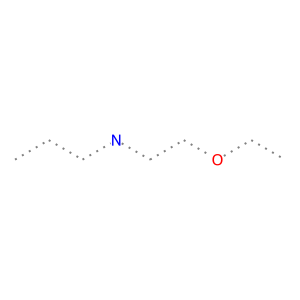 | 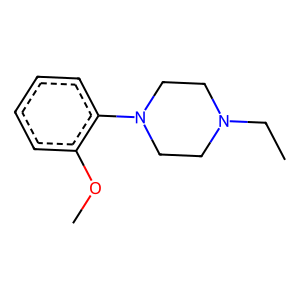 | 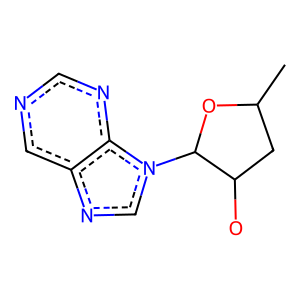 |
| 2 | 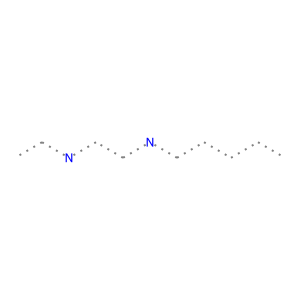 | 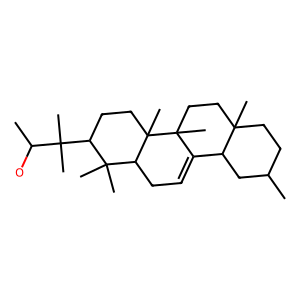 | 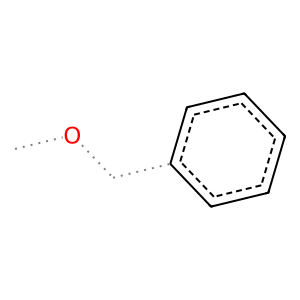 |
| 3 | 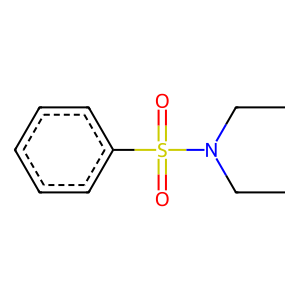 | 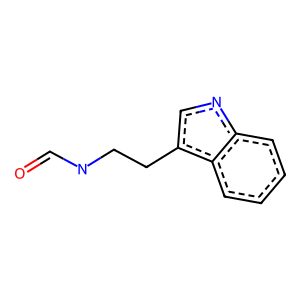 | 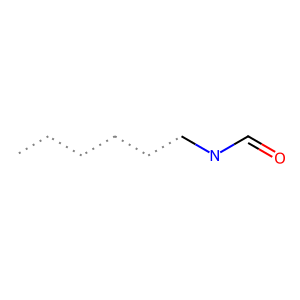 |
| 4 | 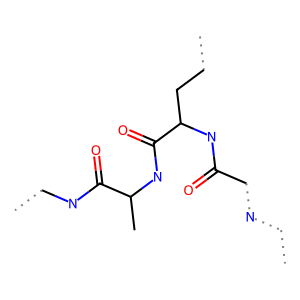 | 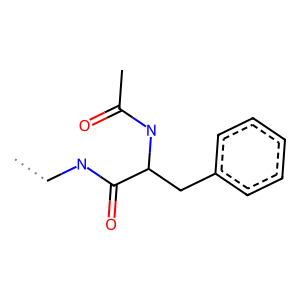 | 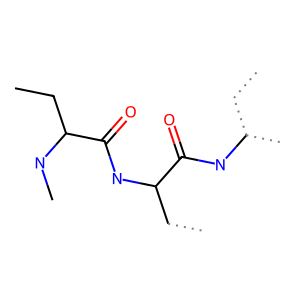 |
| 5 | 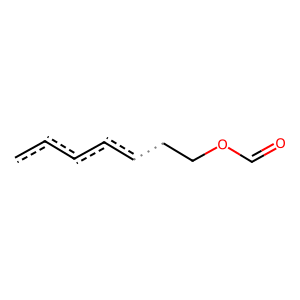 | 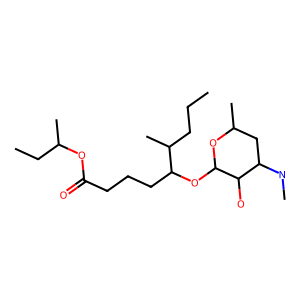 | 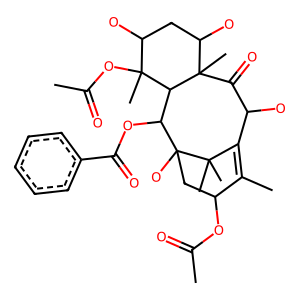 |

**Table S2.** Maximum Common Substructures (MCS) for the five most populated clusters obtained at thresholds of 3, 3.5, and 4 standard deviations above the global iSIM clustered with BitBirch with branching_factor = 1024 and diameter merging criterion for AP fingerprints.

| Cluster | iSIM-sigmas added to iSIM | | |
| --- | --- | --- | --- |
|  | 3 | 3.5 | 4 |
| 1 | 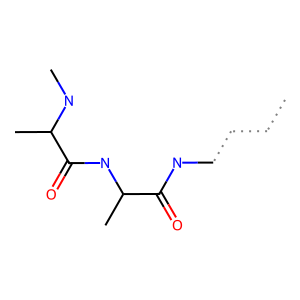 | 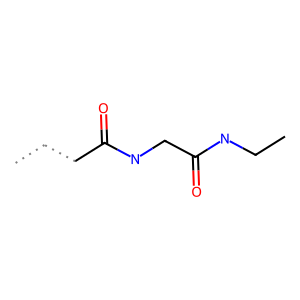 | 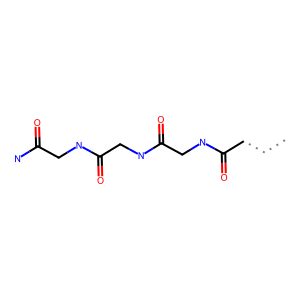 |
| 2 | 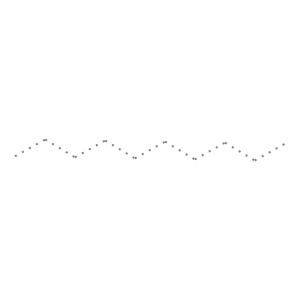 | 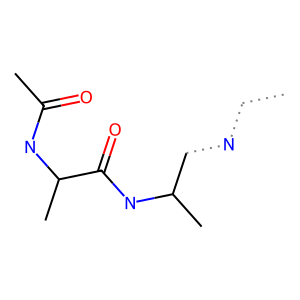 | 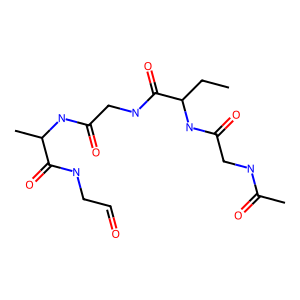 |
| 3 | 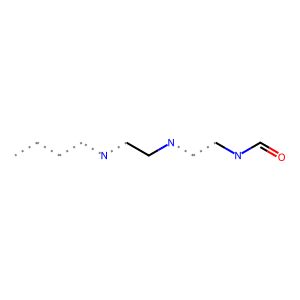 | 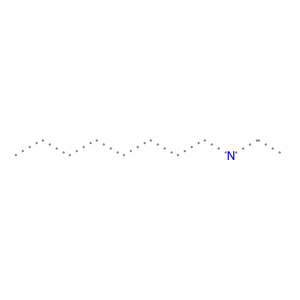 | 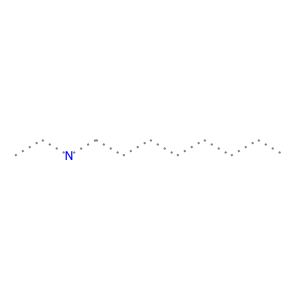 |
| 4 | 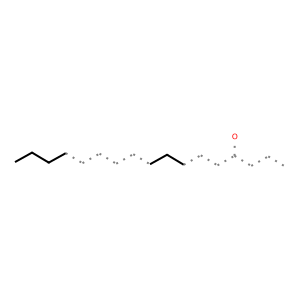 | 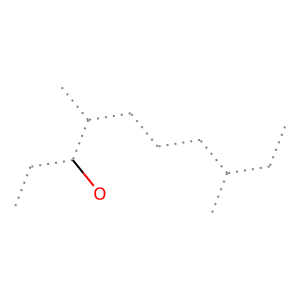 | 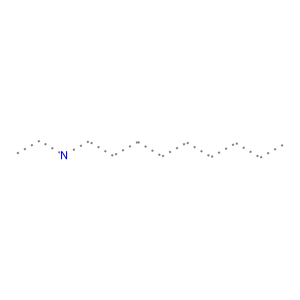 |
| 5 | 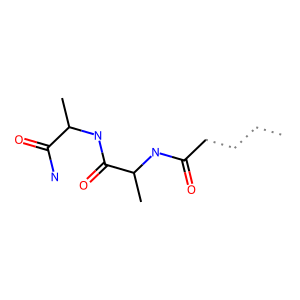 | 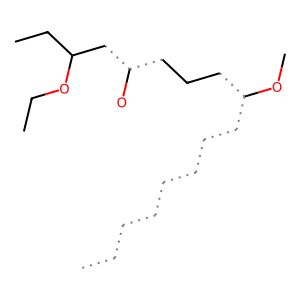 | 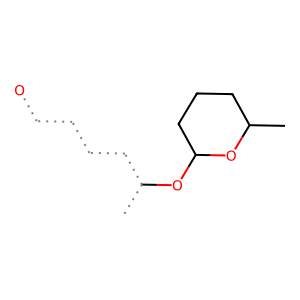 |

**Table S3.** Maximum Common Substructures (MCS) for the five most populated clusters obtained at thresholds of 3, 3.5, and 4 standard deviations above the global iSIM clustered with BitBirch with branching_factor = 1024 and diameter merging criterion for RDKIT fingerprints.

| Cluster | iSIM-sigmas added to iSIM | | |
| --- | --- | --- | --- |
|  | 3 | 3.5 | 4 |
| 1 | 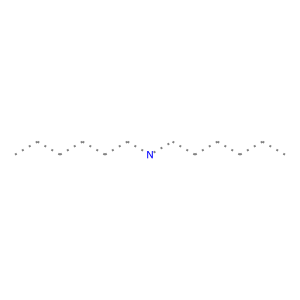 | 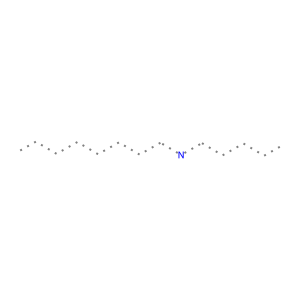 | 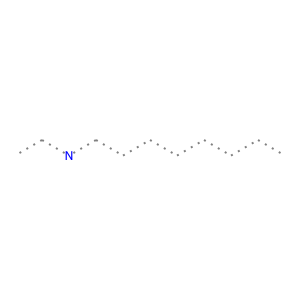 |
| 2 | 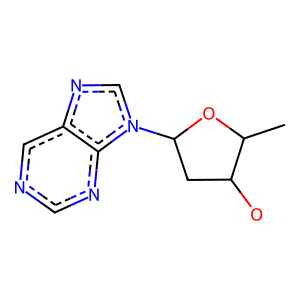 | 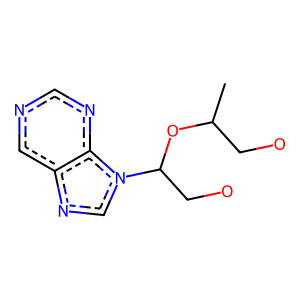 | 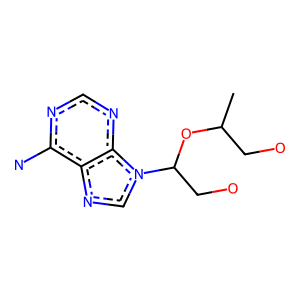 |
| 3 | 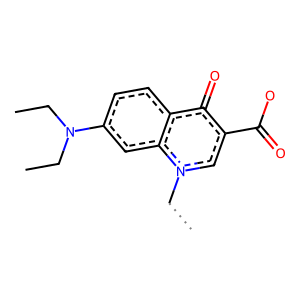 | 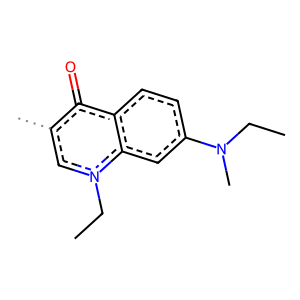 | 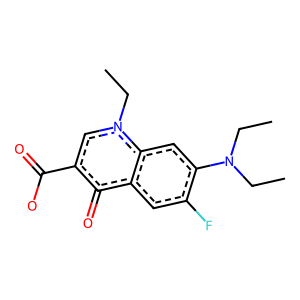 |
| 4 | 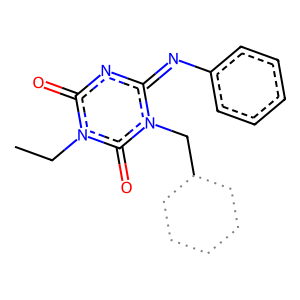 | 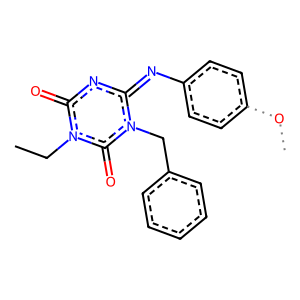 | 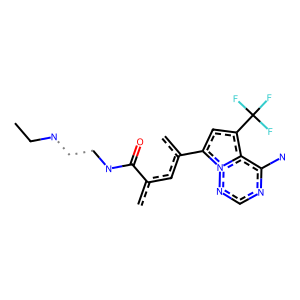 |
| 5 | 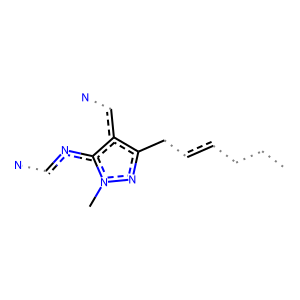 | 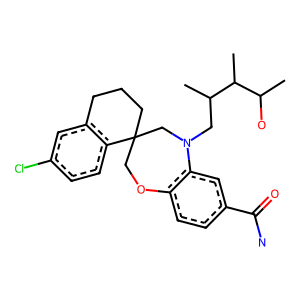 | 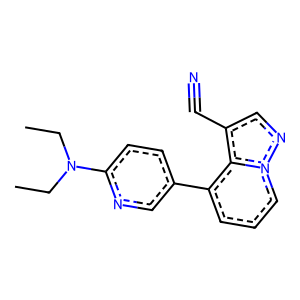 |

**Table S4.** Maximum Common Substructures (MCS) for the five most populated clusters obtained at thresholds of 3, 3.5, and 4 standard deviations above the global iSIM clustered with BitBirch with branching_factor = 1024 and diameter merging criterion for Topological fingerprints.

| Cluster | iSIM-sigmas added to iSIM | | |
| --- | --- | --- | --- |
|  | 3 | 3.5 | 4 |
| 1 | 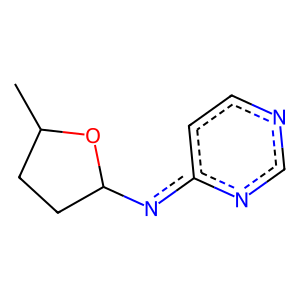 | 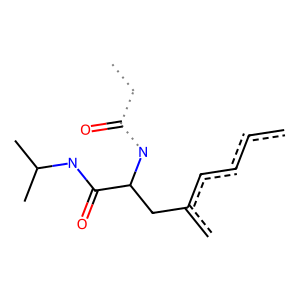 | 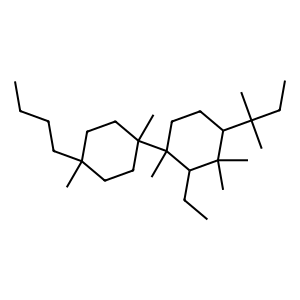 |
| 2 | 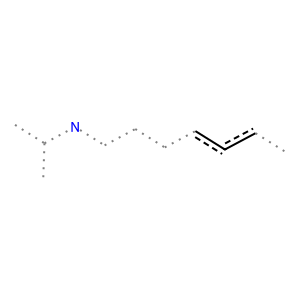 | 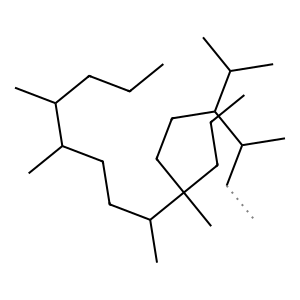 | 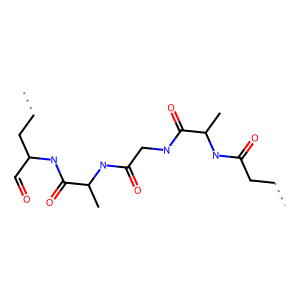 |
| 3 | 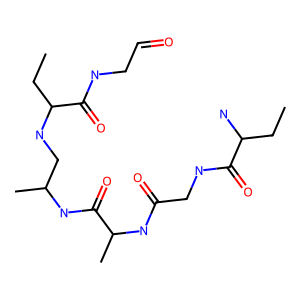 | 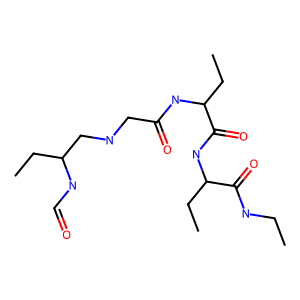 | 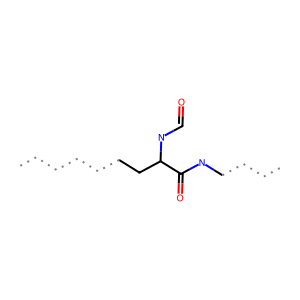 |
| 4 | 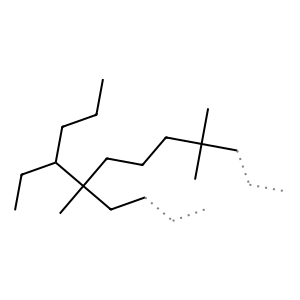 | 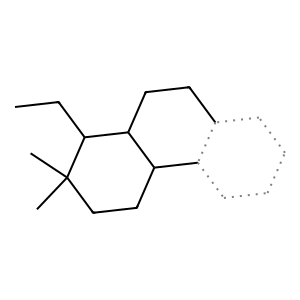 | 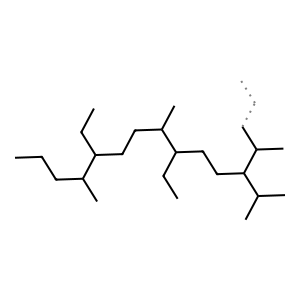 |
| 5 | 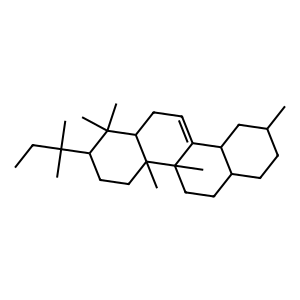 | 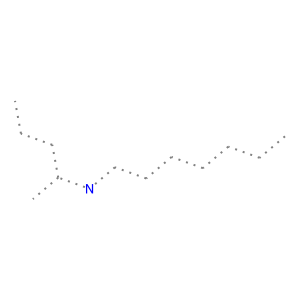 | 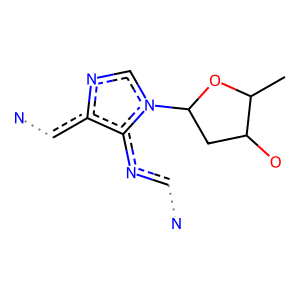 |

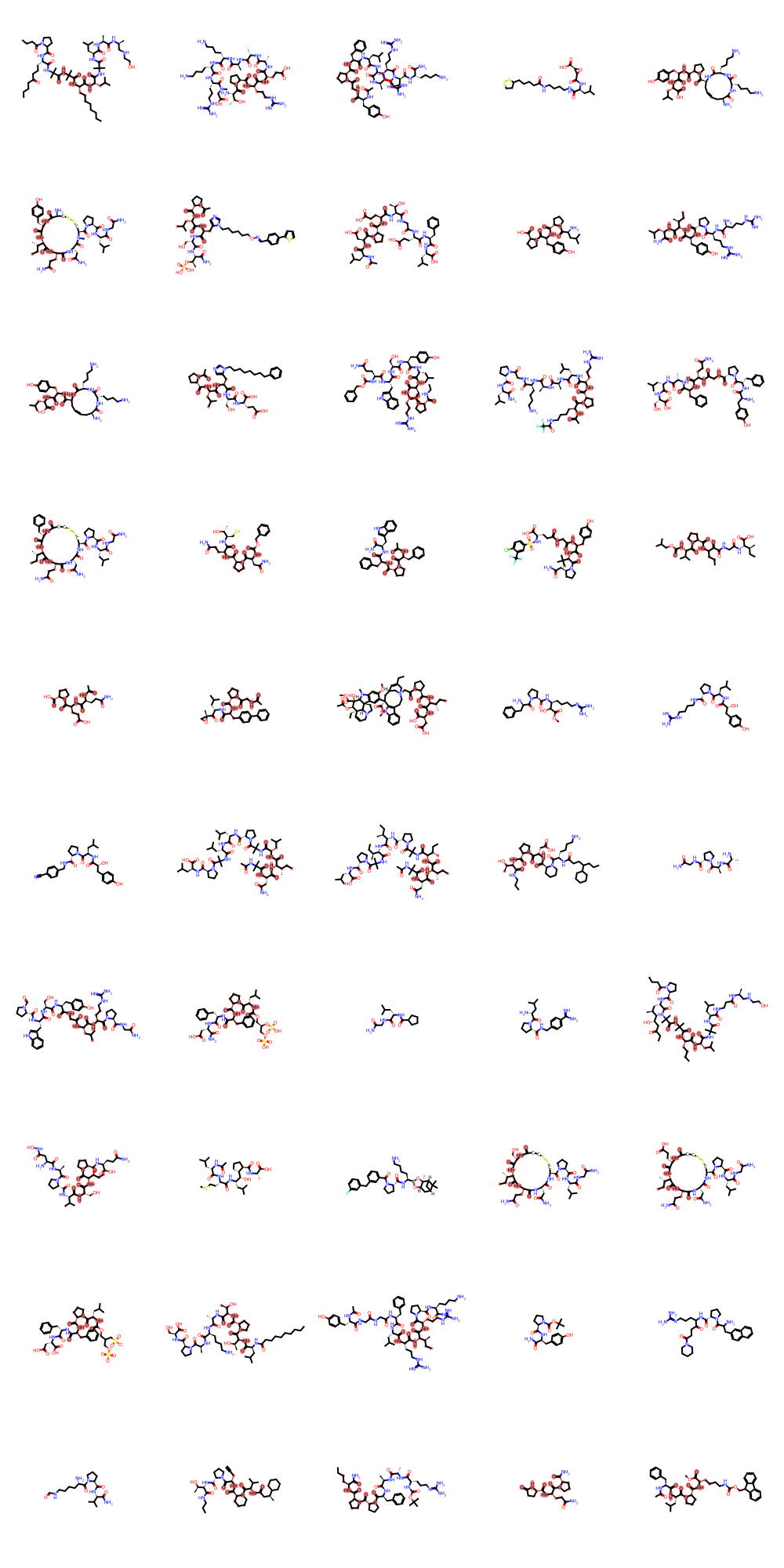

**Figure S11.** Random sampled structures and highlighted MCS for the most populated cluster for ECFP4 fingerprints, threshold of 3 standard deviations over iSIM.

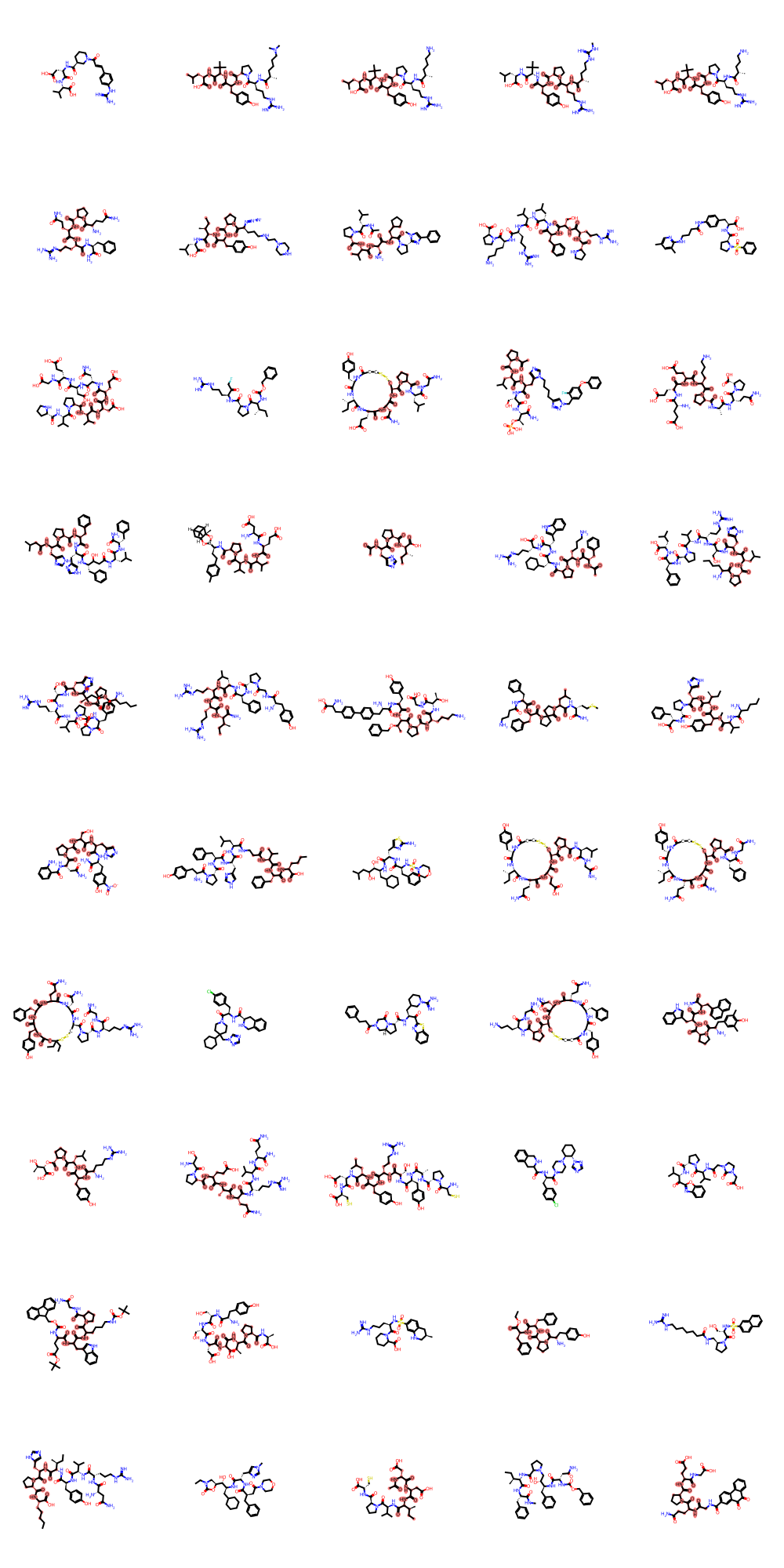

**Figure S12.** Random sampled structures and highlighted MCS for the most populated cluster for ECFP4 fingerprints, threshold of 3.5 standard deviations over iSIM.

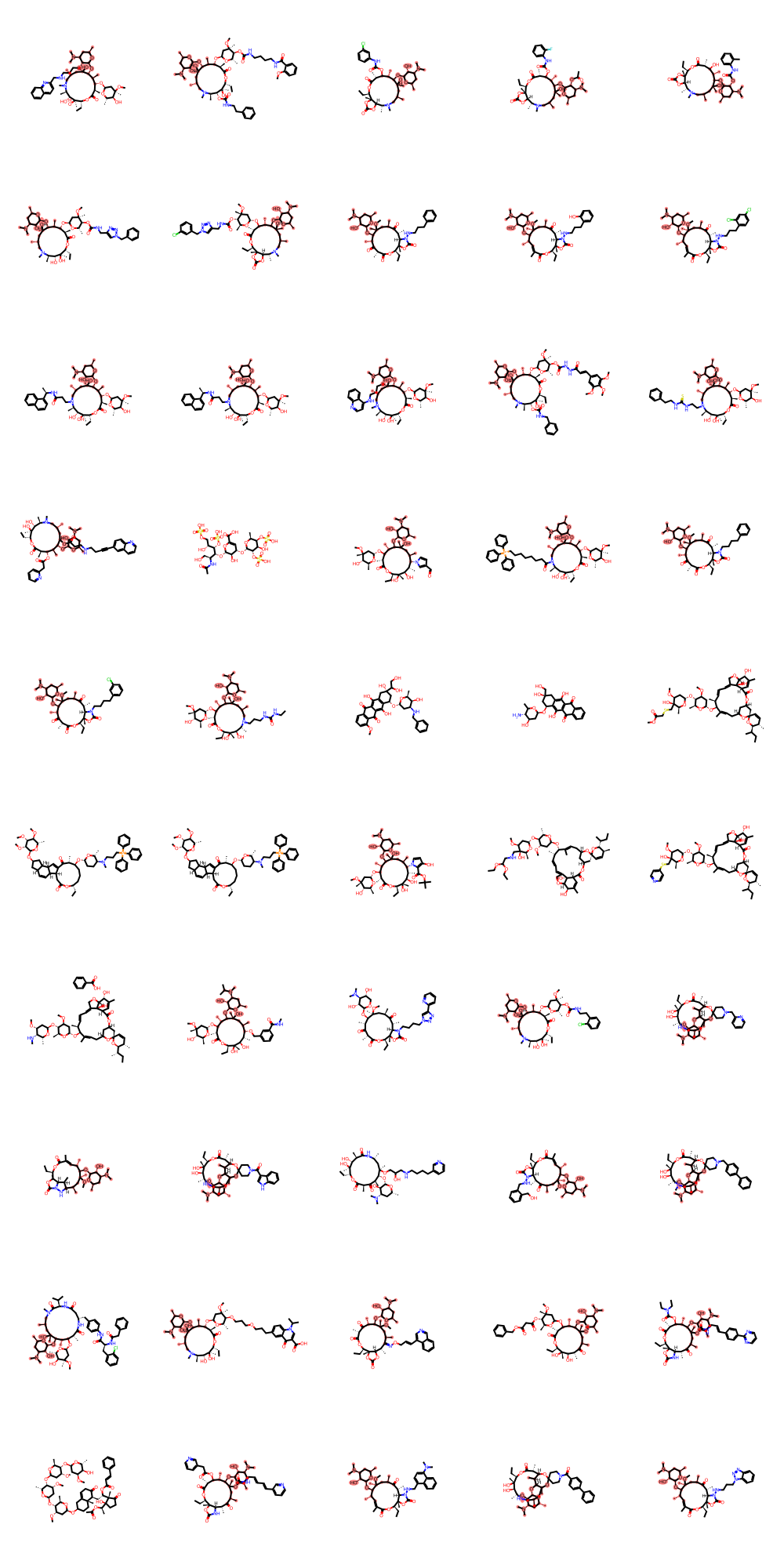

**Figure S13.** Random sampled structures and highlighted MCS for the most populated cluster for ECFP4 fingerprints, threshold of 4 standard deviations over iSIM.

**Table S5.** MCS and populations for the top 20 most populated clusters for the ChEMBL34 library represented with ECFP 2048-bit fingerprints after 5 re-clustering iterations with and without extra-threshold in each step.

| C# | No extra-threshold | 1 σ extra-threshold | C# | No extra-threshold | 1 σ extra-threshold |
| --- | --- | --- | --- | --- | --- |
|  | MCS | MCS |  | MCS | MCS |
| 1 | 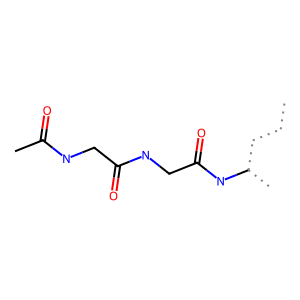 | 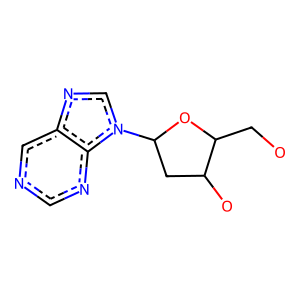 | 7 | 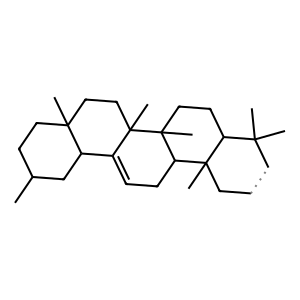 | 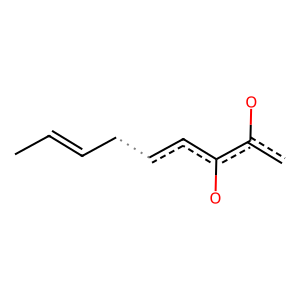 |
| 2 | 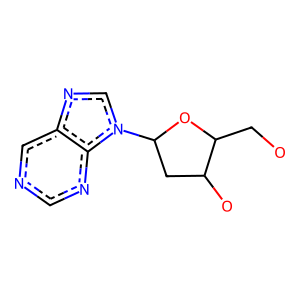 | 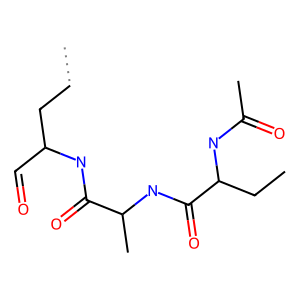 | 8 | 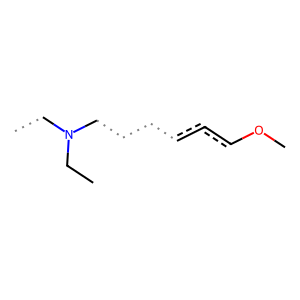 | 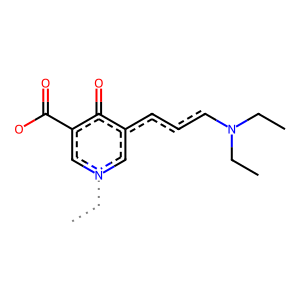 |
| 3 | 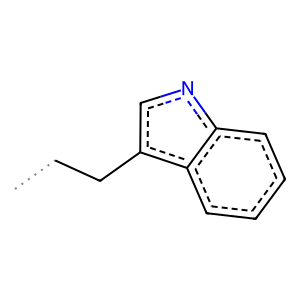 | 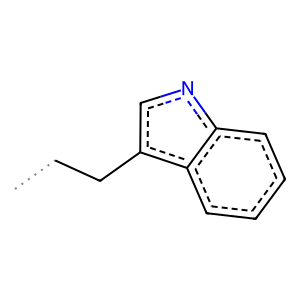 | 9 | 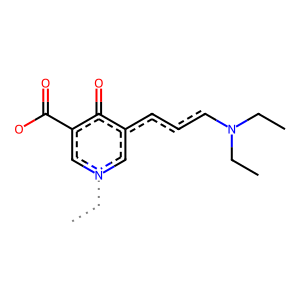 | 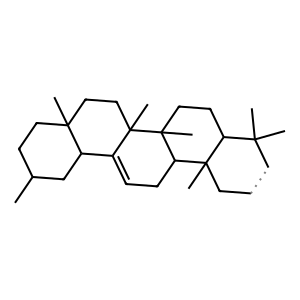 |
| 4 | 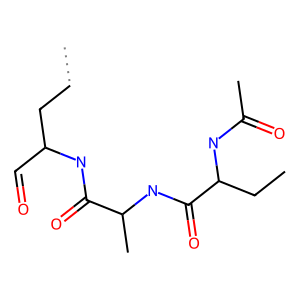 | 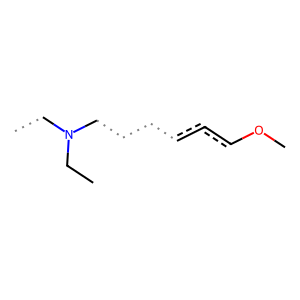 | 10 | 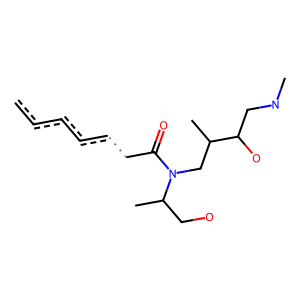 | 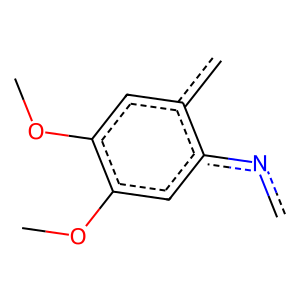 |
| 5 | 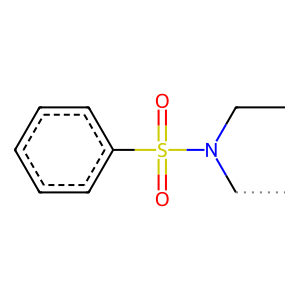 | 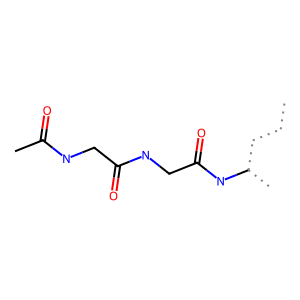 | 11 | 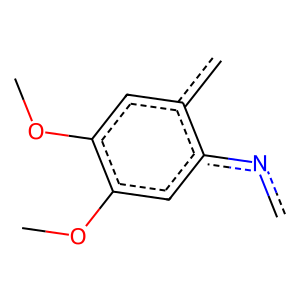 | 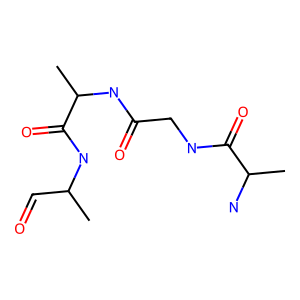 |
| 6 |  |  | 12 |  |  |
| 13 |  |  | 17 |  |  |
| 14 |  |  | 18 |  |  |
| 15 |  |  | 19 |  |  |
| 16 |  |  | 20 |  |  |

Table S6 shows some clusters that were prevented from merging due to the use of an extra-threshold in the re-clustering steps. We can notice how there is a persistence of the MCS core in all the clusters, however the substitution and chemistry of them are different. For instance, we see how cluster 1 has mainly benzyl- and phenyl-alkyl substituents, while cluster 261 has cyclic phosphate derivates, cluster 850 alkyne long chains, and cluster 1191 amide long chains. With this table we want to exemplify that if no extra-threshold is added, different decorations end up in the same cluster, hence it is up to the user necessity if they just want to find clusters with one core MCS, or that are more chemically related in the decorations too. Table S7 shows another example of this.

**Table S6.** Examples of subclusters that are prevented to merge together (yet they have the same MCS) by the use of an extra-threshold for the cluster #2 obtained with no threshold. Indicated cluster numbers correspond to the one standard deviation extra-threshold instance.

Cluster 1

Cluster 261

Cluster 850

Cluster 1191

**Table S7.** Examples of subclusters that are prevented to merge together (yet they have the same MCS) by the use of an extra-threshold for the cluster #7 obtained with no threshold. Indicated cluster numbers correspond to the one standard deviation extra-threshold instance.

Cluster 755

Cluster 281

Cluster 2837

Cluster 196

**Figure S14.** Flow (1<n<100) diagram for the top 20 clusters after 5 re-clustering steps with an extra threshold of 1 standard deviation for ChEMBL34 with ECFP4 fingerprints, branching factor 1024.

**Table S8.** Examples of clusters merged together by doing a re-clustering step with extra-threshold for the most populated cluster (Cluster 0 in Figure S14).

Cluster 14500

Cluster 9284

Cluster 16408

**Table S8.** Examples of clusters merged together by doing a re-clustering step with extra-threshold for a cluster 7 in Figure S14.

Cluster 10958

Cluster 15051

Cluster 21850

Cluster 25070

**Figure S15.** Flow (n=1) diagram for singletons merging into the top 30 clusters after 5 re-clustering steps with an extra threshold of 1 standard deviation for ChEMBL34 with ECFP4 fingerprints, branching factor 1024.

**Table S8.** Examples of singletons merged together by doing a re-clustering step with extra-threshold for the most populated cluster (Cluster 25 in Figure S15).

| Singleton 139245 | Singleton 139317 |
| --- | --- |
| Singleton 139982 | Singleton 140150 |
| Singleton 140187 | |

**Section 2:** Scubidoo library <https://scubidoo.pharmazie.uni-marburg.de/> (*n* = 999,794)

**Figure S16.** Variation of the number of clusters, singletons, singleton-to-cluster ratio; and iSIM of the medoids for the scubidoo library represented with ECFP4 2048-bit fingerprints. iSIM of the global library is indicated with dashed line.

| a) | b) |
| --- | --- |

**Figure S17.** a) Population and unique scaffold distribution for the top 20 largest clusters and b) overall cluster population count for the scubidoo library using BitBirch (branching factor of 1024) and thresholds of 3.5 standard deviations over iSIM.

**Table S9.** MCS for the top 5 clusters of scubidoo library represented by ECFP4 2048-bit fingerprints with using BitBirch (branching factor of 1024) and thresholds of 3.5 standard deviations over iSIM.

| Cluster 1 | Cluster 2 |
| --- | --- |
| Cluster 3 | Cluster 4 |
| Cluster 5 | |

**Figure S18.** Evolution of (a) the number of clusters, number of singletons, and singleton-to-cluster ratio, and (b) the iSIM of the cluster medoids as a function of successive re-clustering iterations. Results correspond to the scubidoo library represented with ECFP4 2048-bit fingerprints and clustered using BitBIRCH with a branching factor of 1024 and a threshold set to 3.5 standard deviations above the global iSIM of the dataset with extra-threshold of one standard deviation.

**Section 3:** COCONUT Carotenoids library (<https://coconut.naturalproducts.net>) (*n* = 1,038)

**Figure S19.** Variation of the number of clusters, singletons, singleton-to-cluster ratio; and iSIM of the medoids for the carotenoid library represented with ECFP4 2048-bit fingerprints. iSIM of the global library is indicated with dashed line.

| a) | b) |
| --- | --- |

**Figure S20.** Population and unique scaffold distribution for the top 20 largest clusters and overall cluster population count for the carotenoids library using BitBirch (branching factor of 1024) and thresholds of 3.5 standard deviations over iSIM.

**Table S10.** MCS for the top 5 clusters of the carotenoids library represented by ECFP4 2048-bit fingerprints with using BitBirch (branching factor of 1024) and thresholds of 3.5 standard deviations over iSIM.

| Cluster 1 | Cluster 2 |
| --- | --- |
| Cluster 3 | Cluster 4 |
| Cluster 5 | |

**Figure S21.** Evolution of (a) the number of clusters, number of singletons, and singleton-to-cluster ratio, and (b) the iSIM of the cluster medoids as a function of successive re-clustering iterations. Results correspond to the carotenoids library represented with ECFP4 2048-bit fingerprints and clustered using BitBIRCH with a branching factor of 1024 and a threshold set to 3.5 standard deviations above the global iSIM of the dataset with extra-threshold of one standard deviation.

**Section 4:** Molport library (<https://www.molport.com/shop/database>) (*n* = 5,287,740)

**Figure S22.** Variation of the number of clusters, singletons, singleton-to-cluster ratio; and iSIM of the medoids for the molport library represented with ECFP4 2048-bit fingerprints. iSIM of the global library is indicated with dashed line.

**Figure S23.** Overall cluster population count for the molport library using BitBirch (branching factor of 1024) and thresholds of 3.5 standard deviations over iSIM before reclustering.

**Figure S24.** Evolution of (a) the number of clusters, number of singletons, and singleton-to-cluster ratio, and (b) the iSIM of the cluster medoids as a function of successive re-clustering iterations. Results correspond to the carotenoids library represented with ECFP4 2048-bit fingerprints and clustered using BitBIRCH with a branching factor of 1024 and a threshold set to 3.5 standard deviations above the global iSIM of the dataset with extra-threshold of one standard deviation.

**Figure S25.** Overall cluster population count for the molport library using BitBirch (branching factor of 1024) and thresholds of 3.5 standard deviations over iSIM after 3 iterations of reclustering with an extrathreshold of one isim sigma.
